# Unsupervised cellular phenotypic hierarchy enables spatial intratumor heterogeneity characterization, recurrence-associated microdomains discovery, and harnesses network biology from hyperplexed in-situ fluorescence images of colorectal carcinoma

**DOI:** 10.1101/2020.10.02.322529

**Authors:** Samantha A. Furman, Andrew M. Stern, Shikhar Uttam, D. Lansing Taylor, Filippo Pullara, S. Chakra Chennubhotla

**Author notes:** Corresponding Author (S. Chakra Chennubhotla), (Andrew M. Stern). New affiliation: UPMC Hillman Cancer Center, Pittsburgh, PA, 15213, USA.

## Abstract

LEAPH is an unsupervised machine **<u>le</u>**arning **<u>a</u>**lgorithm for characterizing in situ **<u>p</u>**henotypic **<u>h</u>**eterogeneity in tissue samples. LEAPH builds a phenotypic hierarchy of cell types, cell states and their spatial configurations. The recursive modeling steps involve determining cell types with low-ranked mixtures of factor analyzers and optimizing cell states with spatial regularization. We applied LEAPH to hyperplexed (51 biomarkers) immunofluorescence images of colorectal carcinoma primary tumors (N=213). LEAPH, combined with pointwise mutual information (PMI), enables the discovery of phenotypically distinct *microdomains*, composed of spatially configured computational phenotypes. LEAPH identified a subset of microdomains visualized as the spatial configuration of recurrence-specific signaling networks whose intracellular and intercellular interactions support cancer stem cell maintenance and immunosuppression in the evolving tumor microenvironment. The LEAPH framework, when combined with microdomain discovery and microdomain-specific network biology, has the potential to provide insights into pathophysiological mechanisms, identify novel drug targets and inform therapeutic strategies for individual patients.

## Introduction

Tumor cells and their stromal cell counterparts that comprise the tumor microenvironment (TME) reciprocally coevolve to generate heterocellular communication networks. A distinctive characteristic of the functional organization of this continuously evolving ecosystem is spatial intratumoral heterogeneity (ITH), a key determinant of disease progression landmarks in multiple carcinomas that include colorectal cancer [1–10]. Therefore, to optimize diagnosis, prognosis, therapeutic strategies and to identify novel therapeutic targets it is important to define spatial ITH in the tumors of individual patients and determine the mechanistic underpinnings of its relationship to metastatic potential, immune evasion, recurrence, therapeutic response and drug resistance. [1, 3, 11, 12].

The first step in investigating spatial ITH is to identify the cell phenotypes within the TME. This, however, is a challenging task owing not only to the diversity of well-defined cell types within the TME but also the intrinsic plasticity of many of these cell types in response to the selection pressures within their particular confines (i.e., microdomains) [13]. Several single cell approaches are well suited to capture this emergent phenotypic continuum, but at the expense of preserving spatial context. Thus, complementary experimental and computational approaches are needed to further define spatial ITH. The choice of construction of a cellular phenotyping algorithm is based on the type of data generated due to the steep tradeoff between cellular resolution, spatial context, and the number of genomic/proteomic probes employed [14–16]. There has been a recent explosion of hyperplexed [17] (> 9 fluorescence or mass spec-based) biomarker labeling and imaging modalities utilizing various reagent technologies to probe the same tissue sections with several dozens of biomarkers at cellular and subcellular resolutions [18–20]. The challenge now is to accurately characterize the complex spatial and high-dimensional output of these hyperplexed techniques.

Single-cell phenotyping analysis exists in various forms, each with advantages and drawbacks. With high-dimensional data, the analysis usually involves dimensionality reduction but observations have been made that cellular subpopulations distinct in multidimensional space overlap visually in the reduced two-dimensional space [21–24]. Common drawbacks of singlecell clustering algorithms include: 1) reliance on supervised methods leading to user-defined cellular phenotypes as opposed to data-driven cellular phenotypes, 2) lack of a spatial context due to the limitations of data generation, 3) use of hard clustering (a cell belongs to one and only one cellular phenotype) which leaves no room to identify cells along a phenotypic continuum. In Table S1, we detail the current state-of-the-art cellular phenotyping methods and analysis studies within the past decade and specify the incorporation of spatial context, soft clustering, hierarchical component, and unsupervised techniques. To our knowledge, there does not exist a cellular phenotyping method and spatial analysis framework that is unsupervised, harnesses the spatial context, and promotes the phenotypic continuum with sensitivity for identifying transitional state cell populations from hyperplexed in-situ fluorescence images.

In this paper, we propose LEAPH, an unsupervised machine **<u>le</u>**arning **<u>a</u>**lgorithm for characterizing cellular **<u>p</u>**henotypic **<u>h</u>**eterogeneity in a tumor microenvironment (Figure 1). LEAPH builds a phenotypic hierarchy of cell types, cell states and their spatial configurations. We applied LEAPH on hyperplexed (51 biomarkers) immunofluorescence images of colorectal carcinoma (CRC) primary tumor tissue samples (N=213). The biomarkers were selected with the interest of sampling a range of CRC and cancer biology TME properties (see Data section).

**Figure 1 -.**
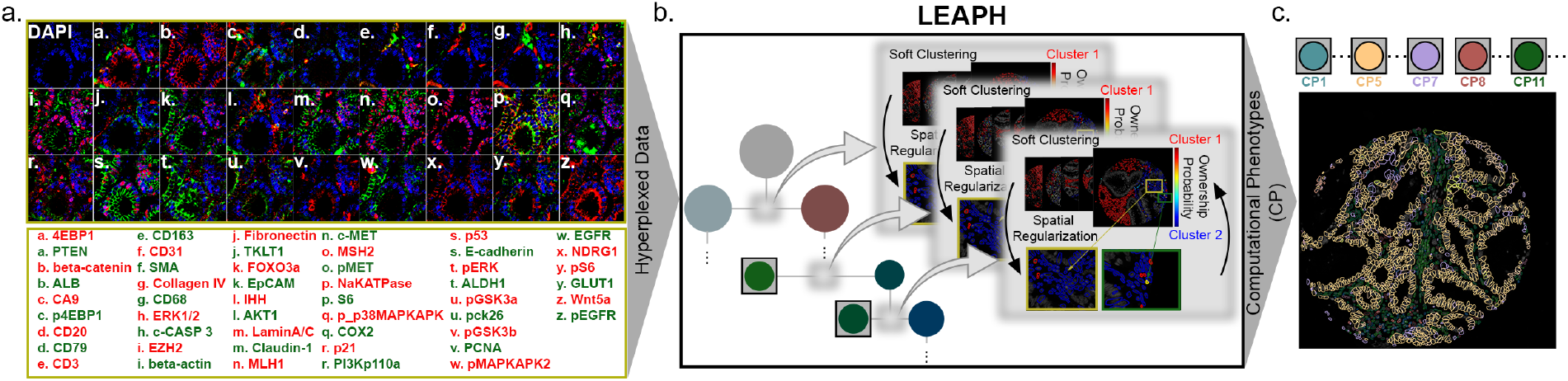
Unsupervised **<u>le</u>**arning **<u>a</u>**lgorithm for cellular **<u>p</u>**henotypic **<u>h</u>**eterogeneity (LEAPH) on hyperplexed colorectal carcinoma tissue microarray dataset. **a.** A zoomed in region of a Stage II CRC tissue samples (ca. 0.6mm cores). Each subpanel is pseudo-colored with DAPI (blue), and pairs of biomarkers indicated in the subpanel (e.g., subpanel a is pseudo-colored with 4EBP1 (red) and PTEN (green)). We selected a large set of biomarkers (p=51) to test the feasibility of cyclical labeling, imaging, and quenching process of the hyperplexing technology and to provide insights into the CRC biology. **b.** LEAPH can perform cellular phenotyping on hyperplexed single sample or a cohort of tissue samples (currently tested up to 213 tissue samples, ~500K cells). LEAPH applies recursive steps of probabilistic clustering with mixtures of factor analyzer (MFA) models and spatial regularization to construct data-driven computational phenotypic trees. We apply a stopping criterion on the recursion, by applying a threshold on the angle between subspaces created by the mixture models, to ensure that the resulting phenotypes are distinct and to avoid overfitting of the input dataset by LEAPH. **c.** The terminal nodes of the tree (leaves) signify distinct computational phenotypes (CP) discovered in the input dataset and form the mixture components of the final MFA model determined by the recursive decomposition. We visualize the spatial distribution of phenotypes within a tissue sample by assigning each cell to the phenotype with the highest ownership probability.

The data-driven and computationally unbiased approach of LEAPH captured a phenotypic hierarchy comprised of *specialized, transitional*, and *multi-transitional* cell states that are intrinsic to the architecturally complex and reciprocally co-evolving TME of CRC. Each LEAPH derived computational phenotype (CP) is further characterized by a unique biomarker signature for ease of interpretation. With these biomarker signatures, we conclude properties demonstrating the heterogeneity of immune cells, cancer stem cells, tumor cells, and cancer associated fibroblasts. In addition, we performed a virtual simulation where the tissue is assumed to be labeled with a subset of key biomarkers (as opposed to the 51 biomarkers we started with) to demonstrate the process to obtain a rank-order subset of biomarkers for characterizing phenotypic diversity.

Using our previous work utilizing pointwise mutual information (PMI) to characterize spatial ITH [25], we show how LEAPH enables the discovery of *microdomains*, which are spatial configurations of computational phenotypes driving disease progression dynamics in the TME of CRC. With a microdomain-specific network biology approach using partial correlation network analysis described in our previous work [26], we find microdomains where the cancer stem cells are driving a Wnt-signaling based immunosuppressive program in patients that will experience recurrence within 3 years post-surgical resection. The LEAPH framework, when combined with microdomain discovery and spatial network biology, has the potential to provide mechanistic insights into disease biology, inform therapeutic strategies for individual patients, identify novel targets for drug discovery and support prognostic/diagnostic predictive analyses.

## Results

### Hyperplexed fluorescence colorectal cancer (CRC) data

The primary source of data for this study, is a cohort of 213 CRC tissue samples (TMA core size = 0.6mm, 1 sample per patient) hyperplexed using Cell DIVE (GE Life Sciences, Issaquah, WA) [18] (Figure 1a). Currently, there exists multiple hyperplexing technologies, each with unique properties. Hyperplex imaging techniques can be non-destructive with the methodology of antibody labeling being either iterative [18] or performed in a batch prior to imaging [19]. There exists other hyperplexing technologies unrelated to fluorescence imaging such as mass cytometry which are both destructive and perform biomarker batch labeling [20].

For this study, our data is imaged using Cell DIVE (GE Life Sciences, Issaquah, WA) [18] which involves non-destructive cyclical immunofluorescence labeling with two or three antibodies labeled with distinct fluorescent probes, followed by imaging and subsequent quenching of the fluorescence. This process is repeated to capture all the required antibodies (biomarkers). The data consists of image stacks taken at each region of interest and overall image stack consists of several images for each of several imaging rounds. Each round includes a nuclear (DAPI) image that is used as a reference for registering all the images from all the rounds. Quantitation of images in each round includes the fluorescence intensity of each measured biomarker. Images are also acquired after quenching rounds for the purpose of autofluorescence removal [18]. Validation of this process shows robustness and preservation of biomarker stability and biological integrity [18, 27–31]. Processing of Cell DIVE images includes correction for uneven illumination across the field of view, removal of autofluorescence, registration, and automated quality control (QC) detection of several categories of defects, including failed registration, blurred or saturated images, and other imaging issues.

The images and data undergo a series of tissue and cell quality checks, log2 transformation and normalization steps. To integrate data from batch processing each biomarker is normalized to a control median. Cellular segmentation is done using a collection of structural biomarkers: NaKATPase (cell membrane, cell border), S6 (cytoplasm), and DAPI (nucleus) (Figure S1). Cells are filtered using individual QC scores generated for each cell (scores less than 0.7-0.8 will not be included indicating inaccurate registration, misalignment, or tissue loss) and based on number of pixels per segmented subcellular compartment. See Methods for more information on data generation and pre-processing steps.

The biomarkers chosen are protein markers for specific cell lineages, oncogenes, tumor suppressors, and post-translational protein modifications indicative of cellular activation states [18] (Figure S2, Table S2). A subset of the biomarkers was selected specifically based on their known association with CRC. In addition to the tissue samples, the CRC patient data set also includes clinical information for each patient regarding sex, age, chemotherapy treatment, and days until recurrence post-surgical resection (patient statistics in Table S3).

### LEAPH builds a phenotypic hierarchy of cell types and states through recursive steps of probabilistic clustering and spatial regularization

LEAPH can process a range of input data from a single sample (~2K cells) to a cohort of samples (tested on 213 tissue samples, ~500K cells total). LEAPH applies recursive steps of modeling cell types with low-ranked mixtures of factor analyzers (MFA) and cell states with spatial regularization to account for the spatial context around each cell (Figure 1b). The *ownership probabilities* for any given cell are constructed recursively by parsing through the hierarchy. Our stopping criterion for recursion is to apply thresholds on the angle between the subspaces created by the mixture models and the fraction of cells with a majority ownership probability in each cluster to ensure that the hyperplexed dataset is not being overfitted (see Methods). The terminal nodes of the tree, i.e., *leaves*, signify distinct data-driven CPs discovered in the input dataset and form the components of the final MFA model determined by the recursive decomposition (Figure 1c).

We define cell states as being *specialized* or *non-specialized* based on the CP ownership probabilities for each cell. A specialized cell state is defined by a strong propensity towards a single CP (ownership probability >0.95). For convenience, we further group the non-specialized cell state as being a *transitional* (ownership probabilities spread between two CPs) or *multi-transitional* (ownership probabilities spread across more than two CPs). We can visualize the spatial distribution of the resulting CP’s within a tissue sample by assigning each cell to the CP with the highest ownership probability (Figure 1c).

To illustrate the process of discovering phenotypic hierarchy, we generated synthetic biomarker data (Figure 2a) to reflect the distribution of biomarker values in the hyperplexed CRC data. In particular, there is a varying spread of biomarker values likely due to the individual biomarker response sensitivity (Figure S4) and the observation of dominant phenotypes (e.g., epithelial and stromal cells) with various nested cell subtypes (e.g., immune, cancer stem cells). To automate the process of phenotypic discovery, we use a recursive probabilistic approach with a low-dimensional latent space mixture of factor analyzers (MFA) (See Methods for more details). Each mixture component is a factor analyzer in a two-dimensional latent space, as we observed this is sufficient to capture the input variance (Figure S5). On the synthetic data (Figure 2a top-left), we instantiate a one-tier MFA model with four mixture components (Figure 2a top-right) and our proposed recursive decomposition where each level of the hierarchy identifies two dominant mixture components (Figure 2a bottom). Comparing the results to the ground truth synthetic data, the one-tier MFA model is unsuccessful at discovering the four phenotypes. The recursive decomposition method separates the larger broad clusters in the first level and the finer sub-clusters at the second level of the hierarchy.

**Figure 2 –.**
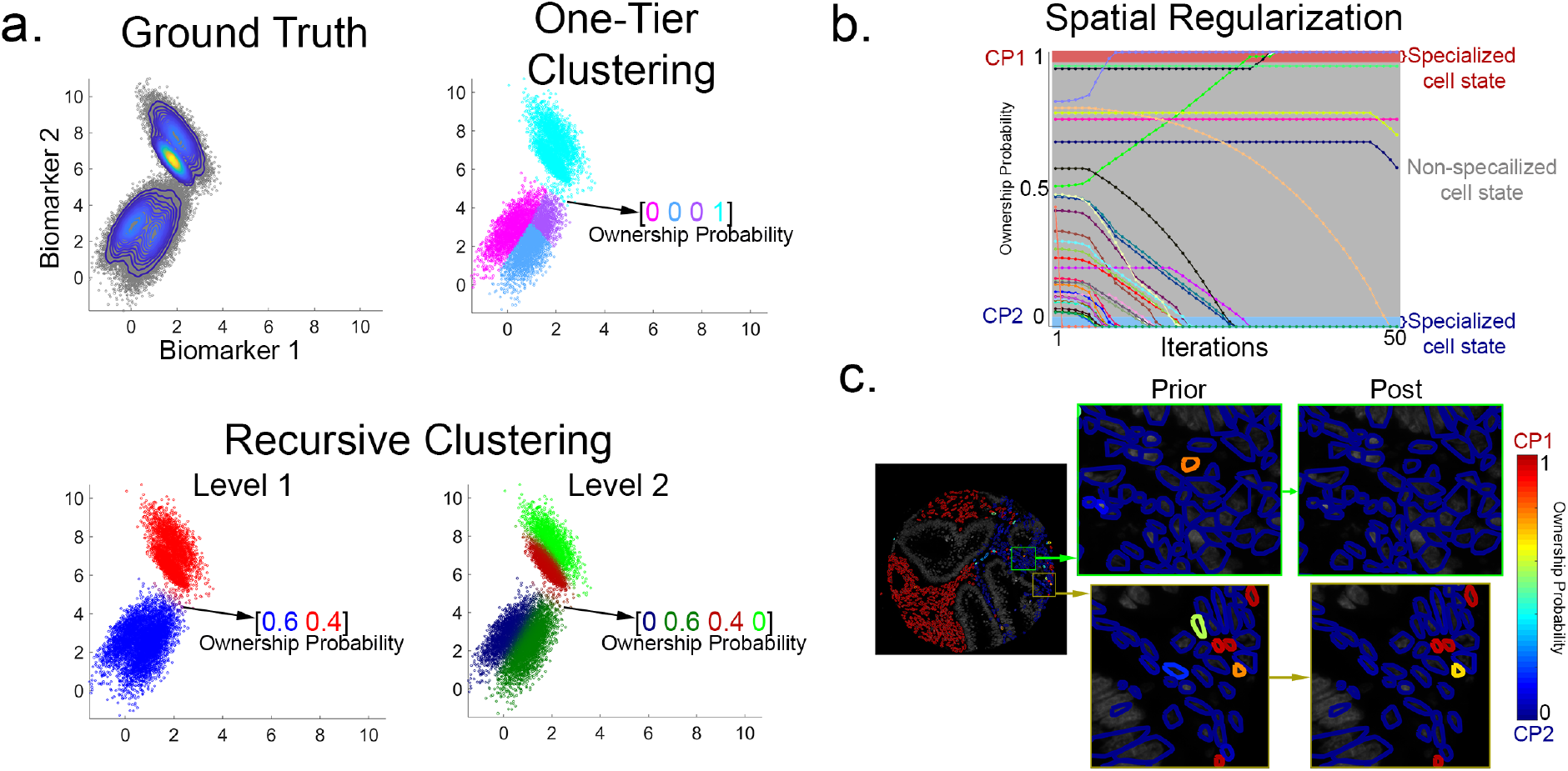
Illustrating the recursive steps of probabilistic clustering with mixtures of factor analyzer models and spatial regularization in LEAPH. **a.** *Top left:* Synthetic data generated to reflect the statistics of the CRC biomarker data. We find that CRC biomarker values have a wide spread likely due to the individual biomarker response sensitivity (e.g., log(ALDH1) values range between 0-12, while log(CD20) values range between 0-4, see Figure S3). Consequently, we observe dominant phenotypes in the form of epithelial and stromal cells and parsing further we discover the various cell subtypes (e.g., immune, STEM cells). To reflect these properties, the synthetic data has two dominant phenotypes each with two additional subtypes and hence four different phenotypes nested in total. ***Top right:*** One tier non-recursive MFA clustering fails at segmenting the four modes in the ground truth synthetic data. ***Bottom left:*** Level 1 of the recursive split identifies the two dominant clusters in the synthetic data. ***Bottom right:*** Level 2 of the recursive clustering splits each dominant cluster from Level 1 to segment the four clusters from the synthetic data. An example ownership probability vector is shown in each clustering attempt, color coded based on respective cluster. This simulation demonstrates the advantage of performing recursive clustering with MFA rather than a one-tier approach. On the more complex and high-dimensional CRC data, each recursive step of fitting MFA model is repeated 100 times and a consensus model is chosen to eliminate any spurious results that arise from the random cluster initialization. **b.** We perform spatial regularization on a single patient sample and track the ownership probabilities for each cell undergoing regularization at each iteration. Majority of cells converge to a specialized cell identity (ownership probabilities >0.95 or <0.05), but a small subset of cells remains in the non-specialized range (ownership probabilities between 0.05 and 0.95) **c.** Tissue sample where each cell is outlined with the ownership probability prior- and post-spatial regularization. We specifically point to two cells, each with an ownership probability of 0.8 prior to spatial regularization. Post-regularization, one cell converges to a specialized CP2 (blue) because of the dominance of this CP in its neighborhood (top zoomed box). The second cell remains non-specialized because of the heterogeneity in its neighborhood (bottom zoomed box).

The MFA model parameters are learned through random initializations over a hundred different runs and a consensus set of model parameters are inferred to capture the optimal subspace representation (Figure S6). The recursive probabilistic clustering when applied along results in a large and unrealistic population of non-specialized cells (typically 25% in a single tissue sample). The soft clustering is agnostic to the spatial complexity of the TME, a key component driving ITH. Based on properties of spatial ITH and the spatial tissue architecture of a tumor, we expect neighborhoods of cells to be spatially coherent (e.g., epithelial/tumor cells to be surrounded by, or spatially proximal to, other epithelial/tumor cells, but making allowance for the presence of tumor-infiltrating lymphocytes and other stromal cells) [32]. To filter false-positive non-specialized cells, we add a spatial regularization component to the recursive probabilistic clustering.

The novel objective function for the spatial regularization component consists of two terms: one to promoting ownership probability confidence and one to promote spatial coherence within a small neighborhood. We assume the two terms hold equal weight and calculate the tuning parameter accordingly. All cells are tested under this objective function but only the cells classified as non-specialized initially (ownership probability 0.05-0.95) are further optimized. Cells initially classified as specialized are fixed (remain invariant during the optimization) to avoid altering the ownerships of cells such as tumor-infiltrating lymphocytes which will *not* maintain spatial coherence. On a single tissue sample, we track the ownership probabilities of the cells undergoing spatial regularization. We observe that most of the cells converge to one cluster, but a subset of cells remains non-specialized (1-4% in a single tissue sample) (Figure 3b). To gain a deeper spatial understanding, we demonstrate the transformation of ownership probabilities of two cells in this simulation. One cell conforms to the surrounding homogenous neighborhood and one cell remains non-specialized between the two clusters due to the heterogeneous nature of the surrounding neighborhood (Figure 3c). A more detailed description of LEAPH can be found in the Methods section.

**Figure 3 –.**
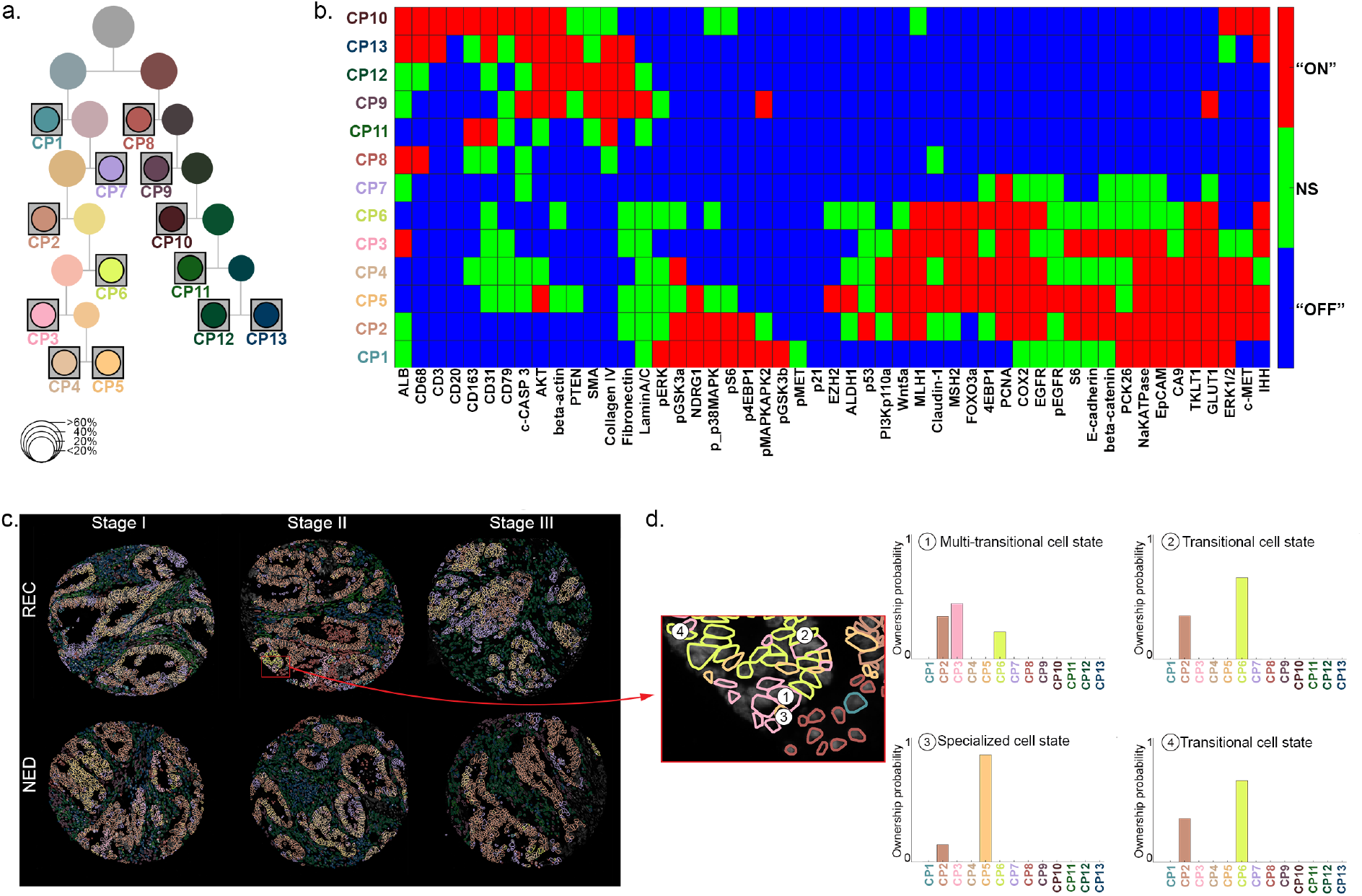
LEAPH reveals the hierarchical nature of the cellular phenotypic heterogeneity and identifies specialized, transitional, and rare cell populations in the hyperplexed colorectal carcinoma dataset. **a.** Cellular phenotypic hierarchy derived by applying LEAPH on the entire CRC cohort. The size of each node is proportional to the fraction of cells with majority ownership to that computational phenotype (CP) (size key). Each leaf node represents a data-driven CP determined by the stopping criteria. The set of all leaf nodes, 13 CPs in total, form the components of the *final* mixture of factor analyzers model for the CRC patient cohort. **b.** For the ease of interpretation, each CP is characterized by a unique biomarker signature, where the biomarkers are classified as being “ON”, “OFF”, or “Non-specific (NS)” (see Methods, Figure S7). The biomarker signatures enable identifying heterogeneous populations of known cell types: epithelial cells (CP6, CP7), tumor cells (CP1, CP2, CP3), cancer stem cells (CP5), cancer associated fibroblasts (CP9, CP12), stromal cells (CP4), macrophages (CP8), and immune cells (CP10, CP13). **c.** Illustrative tissue samples from clinically relevant patient sub-cohorts. Cell boundaries are outlined with the colors of the assigned phenotype (panel a) based on highest ownership probability values. **d.** Example of a neighborhood of non-specialized cells. We point to multiple cells within the neighborhood with varying ownership probabilities. This neighborhood depicts a *multi-transitional* cell state (defined as ownership probabilities spread across more than 2 CPs), *transitional* cell states (ownership probabilities spread across 2 CPs) compared to a specialized cell state (ownership probability greater than 0.95 for a single CP).

### LEAPH captures a diverse set of computational phenotypes having distinct biomarker signatures comprised of specialized, transitional, and multi-transitional cell states and enables the visualization of their spatial configurations

We applied LEAPH on the CRC hyperplexed data (~500K cells) and identified a cellular phenotypic heterogeneity tree consisting of 13 distinct CP’s (leaves) (Figure 3a). We characterize each CP with a unique biomarker signature to describe the intrinsic biological properties (Figure 3b). For each CP, the biomarkers are classified as either “ON”, “OFF”, or “Non-Specific” (“NS”) (see Methods, Figure S7). With the biomarker signatures, we identify a heterogeneous population of epithelial (CP1-7) and stromal (CP8-13) cell types. We can further classify some phenotypes as tumor cells (CP1-3), cancer stem cells (CP5), cancer associated fibroblasts (CP9, CP12), macrophages (CP8), and immune cells (CP10, 13) based on specific indicative biomarkers (Table S2). A small subset of non-specialized cells (7% of ~500K) are dispersed across the 13 CP’s identified.

To visualize the spatial complexity, we assign cells a CP identify based on the highest ownership probability and systematically select representative patients from each outcome-stage based cohort (e.g., NED-Stage I, NED-Stage II,… etc – see Methods for selection process) (Figure 3c). As expected, each tissue sample is comprised of a heterogeneous population of CP’s comprised of specialized, transitional, and multi-transitional cell states (Figures 1c, 3d). In addition to applying LEAPH to the entire patient cohort, we can apply LEAPH to sub-cohorts of patients (particularly based on diagnosis: Stage I, II, III). We do not find a relationship between each stage-specific set of CP’s and recurrence-outcome, but we did notice a difference visually in the spatial distributions. In the next section, we further pursue the spatial relationships between the 13 LEAPH-derived CP’s within each tissue sample to time-to-recurrence and find that the spatial distributions of CP-pairs are statistically significant in relation to time-to-recurrence.

### LEAPH derived computational phenotypes enable the discovery of recurrence-associated microdomains with pointwise mutual information and determine the microdomain-specific network biology

Visually, each tissue sample is composed of a unique pattern of computational phenotypes (from LEAPH). We hypothesize that the patient cohorts with the greatest differences in outcomes will also have distinct spatial heterogeneity patterns. To discover these spatial patterns, we pooled two different cohorts: 1) patients who exhibit recurrence under 3 years (REC-3yrs, N=46) and 2) patients who exhibit no-evidence-of-disease beyond 8 years (NED-8yrs, N=45) (Table S3, see Methods). We use pointwise mutual information (PMI) to quantify the spatial co-occurrence of each CP-pair within each tissue sample. The resulting PMI values are either negative, zero, or positive indicating if a CP-pair spatially co-occurs below, the same, or above average compared to a random background distribution (Figure 4a, see Methods for details).

**Figure 4 –.**
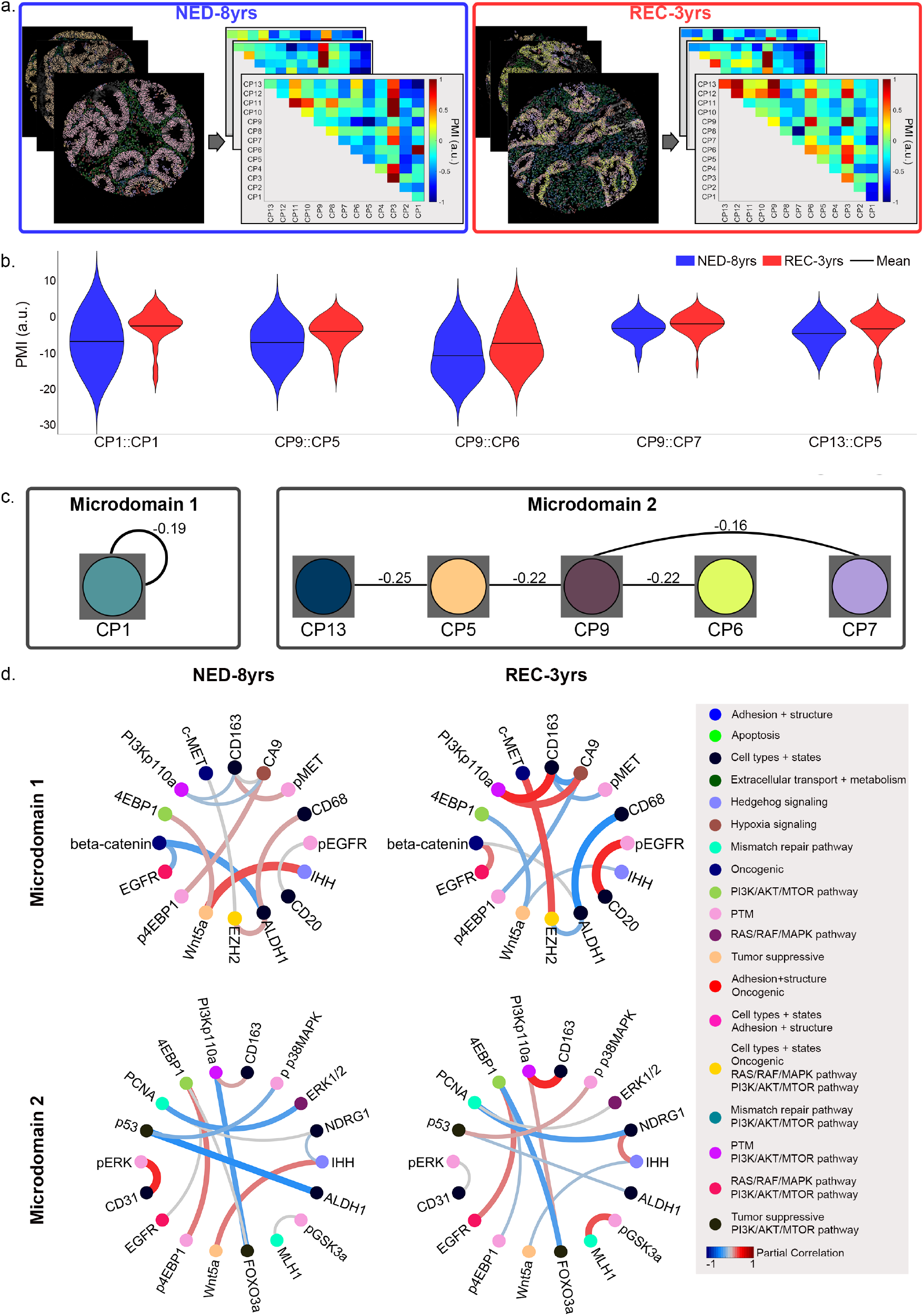
Discovery of recurrence-associated microdomains with pointwise mutual information and determine the microdomain-specific network biology. **a.** Pointwise mutual information (PMI) maps are computed for each tissue sample to quantify the spatial co-occurrence of each CP-pair compared to a random background distribution (see Methods). A PMI value above 0 indicates that the CP-pair spatially co-occurs more often than random and below 0 indicates that the CP-pair spatially co-occurs less often than random. The PMI values are normalized to the range −1 to 1 for visualization. **b.** The PMI values for each CP-pair are grouped by the outcome data: no-evidence of recurrence in 8 years (NED-8yrs) and recurrence within 3 years (REC-3yrs). We compared the PMI distributions using the Kendall rank correlation coefficients and the Wilcoxon rank sum test. We found 5 CP-pairs to be significant (see Methods, Table S4, Figure S9). Each significant CP-pair has a distribution skewed higher for the REC-3yr group implicating a greater level of spatial co-occurrence compared to a random background distribution. **c.** Two microdomains emerge from the significance analysis: microdomain 1 (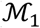 – CP1::CP1) consisting of only a single CP and microdomain 2 (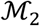 – CP13::CP5::CP9::CP6::CP7) consisting of a network of epithelial and stromal CPs. In the microdomain visualizations, we report the Kendall rank correlation coefficients on each CP-pair edge (p-values for Kendall rank correlation coefficients and Wilcoxon rank sum test are found in Table S4). **d.** Within each microdomain, 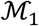 and 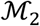, we identify a partial correlation network for the NED-8yrs and REC-3yrs patient cohort consisting of the biomarker pairs with a significant differential in the 99^th^ percentile (see Methods, Table S5 and S6). The biomarkers are grouped based on their presumed cellular functions/processes (Table S2) and the edges connecting each biomarker pair has a color and width corresponding the partial correlation value (also reported in Table S5 and S6).

For each CP-pair, we aggregate the PMI values to form NED-8yrs and REC-3yrs distributions (Figure S9). We use the Wilcoxon rank sum test and Kendall-rank correlations (and p-values) to determine which CP-pairs have a significant difference in spatial co-occurrence between the NED-8yrs and REC-3yrs patients (Table S4, Methods). We report the Kendall-rank correlation between the PMI values and time-to-recurrence in Figure 4b as well as the p-value and results of the Wilcoxon rank sum test in Table S4. In addition, we perform the same statistical analysis using the clinical variables (stage, grade, sex, age) and CP fractions per tissue sample. We found that the spatial distributions (PMI covariates) are statistically more significant in relation to time-to-recurrence than the CP fractions per tissue sample and are similar to the correlation between stage/grade with time-to-recurrence.

We report 5 CP-pairs which spatially co-occur statistically more significantly in the REC-3yrs patient cohort than the NED-8yrs cohort (Figure 4b). The 5 CP-pairs consist of 6 CP’s and form two microdomains (Figure 4c). Microdomain 1 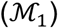 consists of a self-paired CP with heterogeneous properties suggesting the presence of both tumor cells (PCK26+) and tumor associated macrophages (TAMs) (CD163+, Figure S8). The presence of CD163 suggests a protumorigenic M2 polarization [33]. Microdomain 2 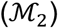 is comprised of an epithelial-stromal network with cancer associated fibroblasts (CAFs) (CP9) as the “hub” interconnecting with an immune cell phenotype (CP13), cancer stem cell phenotype (CP5), and two heterogenous epithelial phenotypes (CP6, CP7) (Figure S8). The association of the stromal cells, particularly CAFs, with poor survival in CRC has been consistently demonstrated [34–36].

We further utilize partial correlation network analysis to investigate the microdomain-specific differential network biology driving the spatial contingency to recurrence outcome (see Methods). The partial correlation measures the true correlation between two biomarkers after the linear dependence from other biomarkers is removed [37, 38]. We perform a connectivity analysis using a permutation test to determine the most significant and differential (99^th^ percentile) biomarker pairwise partial correlations between the NED-8yrs and REC-3yrs cohorts (see Methods, Table S5 and S6) [38]. In Figure 4d, we visualize the outcome-specific differential biomarker pairwise networks, where each edge color and width correspond to the corresponding partial correlation value. Additionally, the biomarkers are color-coordinated based on the properties and functions in Table S2 and Figure S2.

In 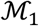, the differential biomarker pairwise partial correlations (Table S5) confirm the presence of cancer stem cells and TAMs supported by Wnt signaling and PI3K/AKT/MTOR in a hypoxic environment as suggested by crosstalk of the later with carbonic anhydrase 9 (CA9). Importantly, each of the partial correlations among the biomarker pairs in 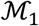 are significantly different when a comparison is made between the NED-8yrs versus the REC-3yrs cohorts. In fact, a majority of these comparisons show a change in sign of the partial correlations suggesting that a distinct difference in network dysregulation in addition to co-occurrence of cell type per se is necessary for driving the recurrence phenotype. The network we have observed for 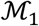 (Figure 4d, Table S5) is consistent with recurrence and the crucial role Wnt signaling plays in cancer stem cell maintenance and tumor cell induced-immunosuppression in CRC and other cancer types [39–47].

Likewise for 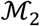, in addition to the aforementioned tumor cell-stromal cell spatial configuration with CAFs as a critical hub (Figure 4c), the difference between the partial correlations among the biomarker pairs in the NED-8yrs versus the REC-3yrs cohorts (Table S6) indicate the critical evolution of network signaling required for driving the recurrence phenotype (Figure 4d).

### Optimal selection of biomarkers for reproducing phenotypic diversity

For hyperplexing platforms that are non-destructive of the tissue and iterative in labeling biomarkers, such as the Cell DIVE platform, there is an option of bringing in biomarkers on demand. In other words, instead of using 51 biomarkers upfront, can we select an optimal list of biomarkers much smaller in number that we can apply iteratively to reveal the phenotypic diversity?

To test this hypothesis, we performed a virtual simulation of systematically introducing biomarkers into the data and applying LEAPH at the first (epithelial-stromal dissection) and second (epithelial- and stromal-subtyping dissection) levels of the hierarchy (see Methods). For comparison, we use the maximum ownership probability from the ALL-DATA results with 51 biomarkers (Figure 3) as the ground truth CP identity. We measure the accuracy as the percentage of cells with a matching CP identity to the ground truth.

At the first level, only 2 biomarkers (pck26, E-cadherin) are needed to reproduce the cellular phenotypic assignments with an accuracy above 97% (Figure S10). Increasing the number of biomarkers further increases the accuracy to almost 99% (Figure S10). This analysis demonstrates that the epithelial-stromal dissection is a low-dimensional dissection, as suspected.

At the second level of the hierarchy, both the epithelial and stromal subtyping dissections require a larger set of biomarkers than level one (Figure S10). The epithelial subtyping dissection reaches 89% accuracy with 4 biomarkers (4EBP1, pS6, pMAPKAPK2, PCNA) (Figure S10). An increase in biomarkers does not make an overall substantial difference in the reproduction accuracy. The stromal subtyping dissection reaches above 93% accuracy with 4 biomarkers (Lamin A/C, Claudin 1, Akt, 4EBP1). Contrasting from the epithelial subtyping dissection, the addition of more biomarkers leads to a convergence to almost 97% accuracy with 8 or more biomarkers (Figure S10).

We conclude that there exists an optimal subset of biomarkers at each LEAPH dissection. This virtual simulation demonstrates the capabilities of LEAPH to exploit the non-destructive and iterative nature of the Cell DIVE platform with a parsimonious selection of biomarkers to regenerate identical computational phenotypes (Figure 5).

**Figure 5 –.**
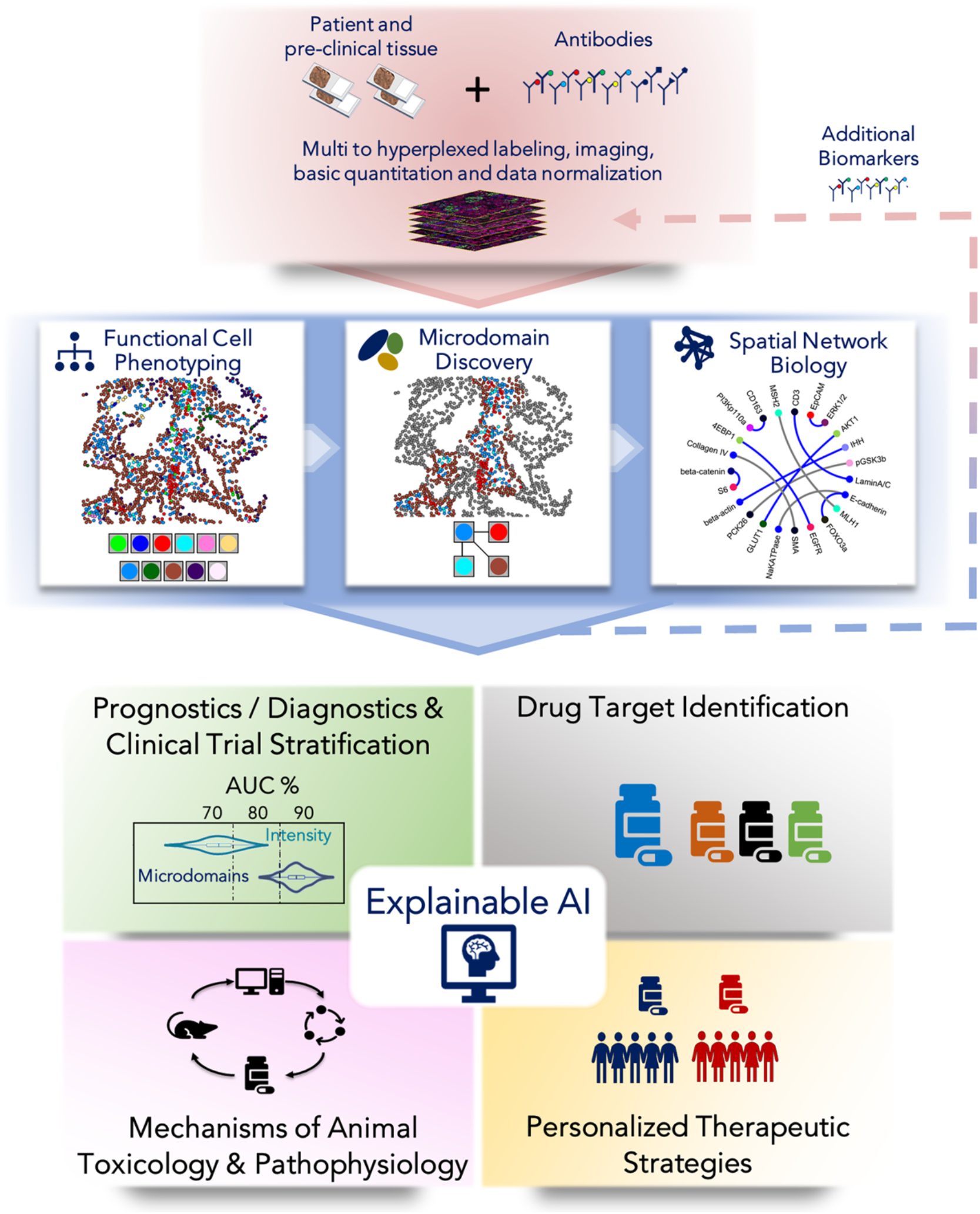
Analytical platform that can be applied to any multi to hyperplexed image data sets. LEAPH enables the application of a spatial analytics and microdomain-specific network biology-based early discovery and development platform to a multi to hyperplexed imaging platform such as Cell DIVE™ used in this study, as well as other hyperplexed technologies [19, 20, 69]. The pipeline begins with the preparation of patient tissue samples for pathology including labeling with multi to hyperplexed biomarkers, imaging based on multiple imaging modalities (e.g. transmitted light, fluorescence, and mass spec [20]). Basic image processing and basic image quantitation is then followed by data normalization in preparation of applying the analytical platform. The hyperplexed imaging process can be initiated with a limited set of biomarkers from which LEAPH builds data-driven, computationally unbiased phenotypic hierarchy of cell types, cell states capturing a phenotypic continuum, and their spatial configurations. LEAPH in combination with pointwise mutual information (PMI) discovers outcome-specific microdomains composed of spatially configured computational phenotypes. LEAPH combined with microdomain-specific network biology provides mechanistic insights into disease biology. This analysis will further suggest outcome-specific new pathways, new cellular phenotypes and additional biomarkers which can be optimally tested through iterative probing of the same microdomains on non-destructive imaging platforms such as Cell DIVE™ [18]. The spatial analytics and microdomain-specific network biology features form the core components of the explainable AI platform [60] that can drive a wide range of applications probing and modulating tumor environment including prognostics, diagnostics, patient stratification for clinical trials, drug target identification, patient-specific therapeutic strategies (e.g. immunotherapy), and animal toxicology studies.

## Discussion

We have proposed LEAPH, an unsupervised, spatially informed, probabilistic recursive clustering method, to exploit the spatial ITH properties prominent in many cancer types and other diseases. The unsupervised and spatially informed approach using biomarker data enables the tumor architecture to drive the discovery of computational phenotypes. Additionally, rather than limiting cells to identify with a single cell type, we exploit the cellular phenotypic continuum with probabilistic clustering and spatial regularization to further classify cell states as specialized or non-specialized. Biologically, the non-specialized cells (transitional and multi-transitional) represent cells undergoing a transformation (e.g., epithelial-mesenchymal-transition, cell-fusion) between CPs. LEAPH does not computationally identify the direction of the transition.

LEAPH is built as a generalizable model amenable to other data sources, soft clustering algorithms, and spatial regularization objective functions/optimization methods. The input criteria for LEAPH is single cell data vectors with corresponding spatial information per sample. Other sources for spatial multiparameter cellular and sub-cellular imaging data include transmitted light (H&E and IHC), fluorescence, immunofluorescence, live cell biomarkers, mass spectrometry, spatial transcriptomics, electron microscopy etc. The soft clustering step of LEAPH is amenable to other probabilistic mixture models such as but not limited to gaussian mixture models [48] and mixtures of probabilistic principal component analysis [49]. An argument could be made that a low-dimensional model will be scalable with a larger number of biomarkers. In addition to the choice of a low-dimensional model, there is room for improvements to better estimate the noise model based on the data generation method. For the spatial regularization step, the objective function in place is amenable to other optimization methods (e.g., gradient descent) and the objective function is adjustable to other sources of input data.

Beyond LEAPH, we have also laid out a framework utilizing the LEAPH derived CPs to investigate the systems biology of TMEs in relation to disease progression. We use PMI to compute the pairwise relative likelihood of spatial co-occurrence between CPs and subsequently form microdomains based on the recurrence-specific CP pairs. We hypothesize that capturing the higher order spatial relationships between groups of phenotypes may provide additional information on the spatial configurations of the microdomains. Further, there are many graphical models [50–52] that could also be used in the microdomain-specific network biology analyses.

In this study, we applied LEAPH on hyperplexed immunofluorescence images of CRC primary tumor tissue samples and identify 13 data-driven CP’s comprised of specialized, transitional, and multi-transitional cell states. We characterized each CP with a biomarker signature and concluded properties demonstrating the heterogeneity of epithelial and stromal cell types. We further used PMI to discover two microdomains with spatial distributions significantly different between patients in the NED-8yrs and REC-3yrs cohorts based on the Kendall rank correlations and Wilcoxon rank sum test.

The two microdomains identified here, 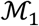 and 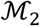 are both more significantly associated with the REC-3yrs cohort than the NED-8yrs cohort. These microdomains comprise spatial configurations of tumor/stromal CPs consistent with those inferred from bulk transcriptomic analyses and resultant consensus molecular signatures that are associated with poor outcome [35, 53]. Nevertheless, these microdomains are also seen in the NED-8yrs cohort suggesting that if the spatial configurations of the CPs within these microdomains are directly involved in recurrence, other factors are also necessary. Importantly, the partial correlation analysis of biomarker pairs showed a strikingly significant difference between the two patient cohorts for both 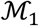 and 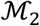. This result supports our working hypothesis that within the evolving tumor microenvironment, the molecular signaling networks within both 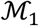 and 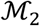 undergo a regulatory switch to confer a recurrence phenotype supported by cancer stem cell maintenance and immunosuppression [39, 40, 42, 44–47, 54–59]. Thus, 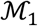 and 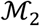 microdomains represent the spatial manifestation of emergent recurrence-specific networks. To further investigate our findings, we can exploit the non-destructive property of Cell DIVE™-like imaging platforms that allow iterative analysis of additional biomarkers on the same tissue sample to test mechanistic hypotheses, identify novel biomarkers and optimal therapeutic strategies (see below and Figure 5).

Here, we denoted the leaves of the tree generated by LEAPH as our CPs. However, information at any level of the LEAPH hierarchy can be used for the spatial analysis described in this paper. In fact, in our previous work [26], we have used only the first level of the LEAPH hierarchy (epithelial, stromal and epi-stroma interface domains) to successfully predict the risk of 5-year recurrence in CRC. Indeed, the same analysis can be repeated by using the entire library of CPs derived from LEAPH. We expect the predictive analysis will be further enhanced with the additional phenotypes and the derived microdomains. Based on the statistical robustness of our approach and external experimental validation of our results found in literature, we predict that we will find similar microdomains presented here when our methods are applied on larger CRC datasets.

The selection of biomarkers is a key factor in determining the resulting phenotypes that will emerge. We demonstrated the existence of an optimal set of biomarkers capable of reproducing the CPs derived from LEAPH with the entire biomarker set. Our virtual simulation provides evidence for performing iterative cycles of imaging and computational analysis with an optimal biomarker set to fully exploit the capabilities of LEAPH in combination with a non-destructive multi to hyperplexed imaging platform.

LEAPH bridges an important knowledge gap in the analytical frameworks that we previously proposed [25, 26]. In particular, LEAPH builds a statistical framework for the application of PMI [25] in the unsupervised discovery of data-driven microdomains. With this advancement, we can fully exploit the spatial network biology based analysis from our previous work [26] to provide mechanistic insights into disease biology such as the generation of outcome-specific new pathways, new cellular phenotypes, and iterative experimental testing of additional biomarkers of the same microdomains (Figure 5). We can generate hypothesis for future experimental and computational studies to further investigate the signaling and cross talk of this immunosuppressive program across the TME landscape to understand the biology of CRC recurrence and possible therapeutic strategies. Systematically deciphering the microdomain-specific network biology will allow us to not only a priori predict recurrence and its aggressiveness resulting in personalized patient surveillance but also potentially select optimal therapeutic interventions by identifying dominant risk-associated microdomains (Figure 5). This framework forms the basis of an explainable AI platform [60] with applications probing and modulating tumor environment including prognostics, diagnostics, patient stratification for clinical trials, drug target identification, patient-specific therapeutic strategies including immunotherapy, as well as animal toxicology studies (Figure 5).

## Acknowledgements

This work was supported in part by the grants NIH/NCI U01CA204836 (SCC, DLT); NIH/NIBIB 5T32EB009403 and 1NIH/NCI 1F31CA254332 (SF); NIH P30CA047904, and PA DHS 4100054875 (DLT); NIH/NHGRI U54HG008540, and UPMC Center for Commercial Applications of Healthcare Data 711077 (SCC). The authors are very grateful to Dr. Fiona Ginty and her team members at GE Global Research, including Dr. Christopher J. Sevinsky, Dr. Yousef Al-Kofahi and Brion Sarachan, for their collaborative support and for sharing the hyperplexed colorectal cancer dataset generated on Cell DIVE™ from their previous study [18].

## Competing Interests

S.C. C. and D.L.T. have ownership interest in SpIntellx, Inc., a computational and systems pathology company. S.C.C is on a leave of absence from the University of Pittsburgh. The other authors disclose no potential conflicts of interest. The computational and systems pathology intellectual property is owned by the University of Pittsburgh and is exclusively licensed to SpIntellx Inc., Pittsburgh, PA.

## Author contributions

Conceived and designed the study – S.A.F., A.M.S., D.L.T., F.P., S.C.C.

Wrote the paper – S.A.F., A.M.S., D.L.T., F.P., S.C.C.

Developed computational platform – S.A.F., F.P., S.C.C.

Performed formal analysis – S.A.F., F.P., S.C.C.

Developed systems biology framework – S.A.F., A.M.S., F.P., S.C.C.

Data quality control and providing analysis support – S.A.F., S.U., F.P., S.C.C.

Technical issues and edited and reviewed the paper – S.A.F., A.M.S., S.U., D.L.T., F.P., S.C.C.

## Methods

### Hyperplexed data

The data for this study consists of 747 colorectal carcinoma (CRC) tissue samples hyperplexed using cell DIVE with 56 biomarkers measured in protein expressions plus DAPI nuclear counterstain. Cell Dive (GE Life Sciences, Issaquah, WA) [18] involves non-destructive cyclical immunofluorescence labeling with two or three antibodies labeled with distinct fluorescent probes, imaging and subsequent quenching of the fluorescence. This process is repeated to capture all the required antibodies (biomarkers). The data consists of image stacks taken at each region of interest and overall image stack consists of several images for each of several imaging rounds. Each round includes a nuclear (DAPI) image that is used as a reference for registering all the images from all the rounds. Quantitation of images in each round includes the fluorescence intensity of each measured biomarker. Images are also acquired after quenching rounds for the purpose of autofluorescence removal [18, 61]. Processing of Cell DIVE images includes correction for uneven illumination across the field of view, removal of autofluorescence, registration, and automated quality control (QC) detection of several categories of defects, including failed registration, blurred or saturated images, and other imaging issues. The images and data undergo a series of tissue and cell quality checks, log2 transformation and normalization steps. To integrate data from batch processing each biomarker is normalized to a control median. Validation of this process shows robustness and preservation of biomarker stability and biological integrity [18, 27–31]. Images are acquired in TIFF format, while image metadata is captured in files having a simple structure that captures the provenance of which images were derived from which slides and characteristics of the acquisition [18, 61]. The biomarkers chosen are protein markers for specific cell lineages, oncogenes, tumor suppressors, and post-translational protein modifications indicative of cellular activation states (Table S2, Figure S2). The data also includes corresponding clinical information including the histological tumor grade, cancer stage, gender, age, and follow up monitoring for 10 years (Table S3).

### Data pre-processing

#### Cell quantitation

Cellular segmentation is done using a collection of structural biomarkers: NaKATPase (cell membrane, border), S6 (cytoplasm), and DAPI (nucleus) (Figure S1). Protein expression and standard deviation were quantified by the median biomarker intensity value within each cell mask and transformed to the log2 scale [18]. Cells are filtered using individual QC scores generated for each cell (scores less than 0.7-0.8 will not be included indicating inaccurate registration, misalignment, or tissue loss) and based on number of pixels per segmented subcellular compartment.

#### Patient selection

For this study, based on the clinical data, we limit the patient data set to deceased patients with recurrence within 5 years (postsurgical resection) and alive patients with no evidence of recurrence within 5 years (post-surgical resection). Further, we eliminate tissue samples with less than a threshold of 1000 cells to limit the potential adverse effects of hyperplex imaging (i.e. damaged, folded, or lost tissue). This cell threshold is computed based on the 20th percentile of number of cells per tissue sample shown in Figure S3. The final data set used is composed of 213 TMA spots (Table S3). The alternative strategies could have been to filter out cells within the damaged areas of the TMA’s by examining the nuclear-to-cytosolic ratio of structural biomarkers. We chose to use the cell threshold method to preserve the unsupervised nature of LEAPH.

#### Biomarker selection and distributions

We removed biomarkers showing batch affects resulting in a selection of 51 biomarkers for this analysis. The distribution of each biomarker (log2 scale) across all cells in the patient cohort is shown in Figure S4. The Kurtosis values measures the skewness of the distribution. Comparing the kurtosis values of the biomarker distributions to the kurtosis of a univariate normal distribution (kurtosis = 3), many but not all these distributions can be considered to have a Gaussian shape.

#### LEAPH: unsupervised learning algorithm for cellular phenotypic heterogeneity

We will describe the hyperplexed dataset in a high-dimensional space, where each cell 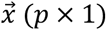 is described by a *p* dimensional vector of biomarker expressions quantitated appropriately. Further, we assume that the hyperplexed dataset has an intrinsic low-dimensional representation. We will use a mixture of factor analyzers described by low-dimensional factor loadings (Λ (*p* × *k*)), latent variables 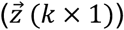, mean vector 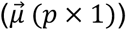, and noise term 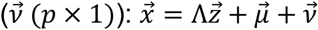, where *p* is the number of biomarkers and *k* is the low-dimension latent space [62]. The latent factors, 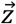, are generated from zero-mean, unit-variance Normal distribution *N*(0, I), and the noise term, 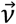 is sampled from *N*(0, Ψ). I is the unit variance and Ψ is assumed to be a diagonal matrix. With this construction, 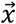 is distributed with zero mean and covariance ΛΛ^*T*^ + Ψ [62].

#### Soft clustering

Typically, cellular phenotyping methods are constructed under the assumption that each cell belongs to one and only one cluster (hard clustering) leaving no room to identify specific cells that may belong to more than one phenotype due to an existing phenotypic continuum. With a Mixture of Factor Analyzers (MFA), we model the cells as *M* components (clusters) with the parameters 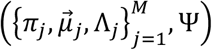 where *π_j_* is the component weight: 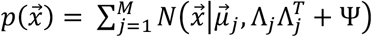. We chose a two-dimensional latent space for each component in the MFA model, as we observed this is enough to capture the input variance (Figure S5). The expectation-minimization (EM) algorithm is utilized to estimate the model parameters [62]. The EM algorithm is initialized with a random set of parameters and the EM algorithm is not guaranteed to converge to a globally optimal solution. To account for this and ensure stability, we perform a hundred different EM optimizations, each initialized randomly. Each optimization yields an MFA model with a set of model parameters. We compute the biomarker ranking for each set of model parameters (see discriminative biomarkers section) and aggregate all biomarker rankings to compute their mean ranking. The model with a biomarker ranking closest (Euclidean distance) to the mean ranking is selected as the consensus model and deemed to provide an optimal subspace representation (Figure S6). The MFA model results in soft clustering probabilities (*ownership probabilities*) – each cell, *x_c_*, holds a unique probability of belonging to each cluster *j*, denoted as Ω_*cj*_.

#### Spatial regularization

The soft clustering is agnostic to the spatial complexity of the TME, a key component driving ITH. Based on properties of spatial ITH and the spatial tissue architecture of a tumor, we expect neighborhoods of cells to be spatially coherent (e.g., epithelial/tumor cells to be surrounded by, or spatially proximal to, other epithelial/tumor cells, but making allowance for the presence of tumor-infiltrating lymphocytes and other stromal cells for example). To promote specialization in cells, we add a spatial regularization component to optimize the ownership probabilities of nonspecialized cells. The spatial regularization step optimizes the objective function which consists of two terms: ownership confidence and spatial coherence given by 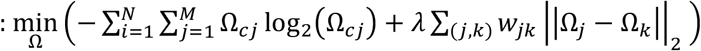. The first term minimizes the entropy of the ownership probabilities promoting specialization in cells. The second term promotes spatial coherence where *w_jk_* is the weight between cell *i* and cell *j* and is computed as the reciprocal of the distance between two cells: 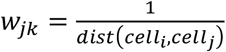. We use a distance threshold (100 pixels at 0.5 *μ*m/pixel) to eliminate an influence between cells that are too far apart to communicate [63].

The objective function is optimized using Alternating Directions Method of Multipliers (ADMM) [64]. We assume that the probability ownership confidence (term 1) and spatial coherence (term 2) should hold equal weight and therefore compute the tuning parameter, *λ*, to scale term 2 to the range of term 1: 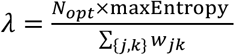 where *N_opt_* is the number of cells being optimized and maxEntropy is the maximum value of the entropy function (=1). Relaxing the assumption that spatial coherence and ownership confidence should hold equal weight in the objective function would lead to a larger parameter space. A higher weight for spatial coherence results in homogenous neighborhoods and a larger set of non-specialized cells. On the contrary, a larger weight for ownership confidence results in the abolishment of all non-specialized cells. We have found stable and consistent results when the tuning parameter represents an equal weighting. Cells can only have neighbors within the same tissue sample and therefore to increase computational speed and efficiency, spatial regularization is performed on each tissue sample independently.

#### Recursive decomposition

To automate the process of phenotypic discovery, we propose a recursive probabilistic approach where each step dissects the most dominant clusters with *M* = 2 components. At each recursive step, the soft clustering step utilizes a low-dimensional latent space MFA. Subsequently, spatial regularization optimizes the resulting per-cell ownership probabilities to filter false-positive non-specialized cells by promoting ownership confidence and spatial coherence. The resulting parameters (ownership probabilities, Ω_*j*_, mean vector, 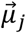, factor loadings, Λ_*j*_) for each cluster, *j*, are passed to the next recursive step to decompose each cluster into further sub-clusters. This process is continued until an attempted cluster split invalidates any one of the following stopping criteria: 1) a resulting cluster takes ownership of < 1% of cells, or 2) the angle between the mean vectors and factor loading space are both below a given threshold.

#### Discriminative biomarker order

Each LEAPH split results in two clusters with high dimensional mean vectors 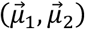. To determine the discriminative ordering of the biomarkers, we compute and sort the proportional difference for each biomarker *j*: 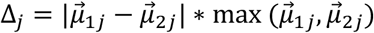. The absolute difference of the mean vectors may bias the selection of biomarkers with high biomarker value ranges and therefore, we opt for a proportional difference to place the biomarkers on an even level for comparison.

#### Interpreting Computational Phenotypes

We characterize each CP with a unique biomarker signature for ease of interpretation. For each biomarker and CP pair two histograms are computed: 1) histogram of biomarker values for all cells (*h_A_*) and 2) histogram of biomarker values for all specialized cells in the CP (*h_B_*). We compare the 20^th^ and 80^th^ percentiles of each histogram to determine the CP-biomarker classification. Each classification is as follows:

| Classification | Operator | Interpretation |
| --- | --- | --- |
| “ON” | $h_B^{80th} > h_A^{80th}$ AND $h_B^{20th} > h_A^{20th}$ | $h_B$ is skewed to the right compared to $h_A$ |
| “OFF” | $h_B^{20th} < h_A^{20th}$ AND $h_B^{80th} < h_A^{80th}$ | $h_B$ is skewed to the left compared to $h_A$ |
| “NS” (Non-specific) | $h_B^{80th} > h_A^{80th}$ AND $h_B^{20th} < h_A^{20th}$<br>OR<br>$h_B^{80th} < h_A^{80th}$ AND $h_B^{20th} > h_A^{20th}$ | $h_B$ spans across $h_A$ or is confined within $h_A$ |

Figure S7 depicts each distribution and 20^th^/80^th^ percentiles for each CP-biomarker pair to demonstrate how the biomarkers are classified in Figure 3b.

#### Selecting representative patients for visualizing computational phenotypes

Post LEAPH, we assign cells a CP identity based on the highest ownership probability value. For *C* CP’s, we compute the fraction of each phenotype found in each patient, *i*, forming a vector 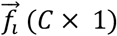. When computing this fraction, we only consider the specialized cells (ownership probability >0.95) to avoid the transitional and rare cells biasing our results. For each outcome-stage-based group (e.g., NED-Stage I, NED-Stage II, NED-Stage III), we compute the average phenotype fraction vector, 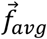, and choose the patient closest to the mean to display in Figure 3c (Euclidean distance).

#### Biomarker selection virtual simulation

For comparison, we assume the LEAPH derived CP’s is the ground truth. Based on the rank-ordered biomarkers from each LEAPH recursion step, we re-run LEAPH with the first 2, 4,…, 20 biomarkers at the first and second level of the hierarchy (Table S7). Assigning each cell to one CP based on the highest ownership probability (cell-label), we compute the accuracy of the cell-labels derived from each sub-set of biomarkers to the assumed ground truth (Figure S10). This virtual simulation demonstrates the reproducibility and capabilities to integrate LEAPH into an integrated iterative imaging and computational platform.

### Discover recurrence-associated microdomains with pointwise mutual information and determine microdomain-specific network biology

#### Step 1: Select patient sub-cohorts with recurrence under 3 years and with no evidence of recurrence in 8 years

To study the differences between patients at each extremum and balance the NED/REC cohorts, we prune the patient cohort further based on the time to recurrence. We consider a group of patients with no evidence of disease over 8 years (NED-8yrs, N=45) and a group of patients that exhibit recurrence under 3 years (REC-3yrs, N=46). See Table S3 for the patient statistics.

#### Step 2: Construct a spatial network of interacting cells in each patient sample

In each tissue sample, we build a spatial network where each node is a cell and the edges connect cells, say *m* and *n*, to each other with a weight *w_mn_* = 1 if their spatial distance *d_mn_* is within a threshold (100 pixels/50 *μ*m, which is the same threshold used in LEAPH spatial regularization), or *w_mn_* = 0 otherwise. To remove possible artifacts, we prune each network to include only cells within the largest connected component.

#### Step 3: Use Pointwise Mutual Information (PMI) to quantify spatial co-occurrence between CP-pairs

Pairwise association statistics, specifically pointwise mutual information (PMI), identifies microdomains consisting of one, two, or groups of phenotypes which spatially co-occur more frequently than a given background distribution. PMI maps can characterize associations at the cohort, patient, and tissue sample level and can describe both the local and global spatial heterogeneity scenarios that cannot be captured by well-known methods such as quadratic entropy [65].

Each cell, *c*, holds an ownership probability vector over the *M* CP’s, 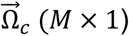, such that 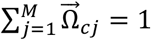. The pointwise mutual information between two phenotypes, (*f_i_, f_j_*), for a given network or network set, *s*, is defined as:

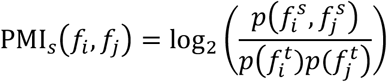

where 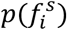 is the probability of phenotype *i* occurring in a network set *s* and 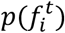 is the background probability distribution of phenotype *i* (this can be an ensemble of networks or individual networks, see below). For a single patient, the joint probability is computed as:

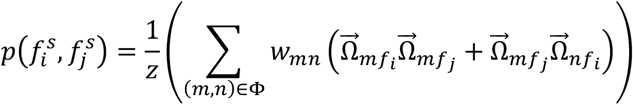

where *ϕ* is the set of edges, *w_mn_* is the edge weight between cells *m* and *n*, and *Z* is the normalization factor given by:

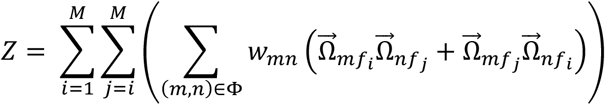

PMI computation results in values with direct implication to spatial co-occurrence. The PMI values are either negative implicating that a phenotype-pair spatially co-occurs less than the background distribution, positive implicating that a phenotype-pair spatially co-occurs more than the background distribution, or zero implicating that a phenotype-pair spatially co-occurs the same as the background distribution.

#### Step 3a: Select a background distribution

We choose to use a random background distribution to depict the probability each CP-pair spatially co-occurs more, less, or the same as random. To construct the random distribution, we set the probability of each CP to the probability of each CP over all cells: 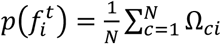, where *N* is the total number of cells.

#### Step 3b: Perform Jackknife estimation

A jackknife estimation is commonly used to remove bias when measuring the dependence between two random variables (e.g., Pearson’s correlation coefficient, mutual information) [66–68]. For a given estimator, PMI: *T_n_* = *T*(*X*_1_,…, *X_n_*) for *n* samples. Let *T*_(−*i*)_ denote the statistic where the *i*-th patient is removed. The jackknife bias estimate is defined as: 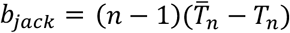, where 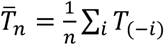. The bias-corrected estimator is

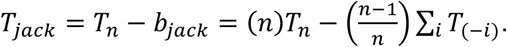

The jackknife estimate of standard error is: 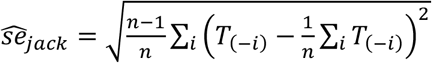.

#### Step 3c: Filter unstable PMI values

We observed instability in the PMI estimates (including high jackknife estimated standard error) for any given CP-pair, (*f_i_, f_j_*) when the number of cells involved in the pairing is low. To remove this bias, we compute the effective number of cells (total ownership probability) in the CP-pair for a patient, *p*, as the summation of ownership probabilities, 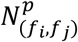, over all cells in the network: 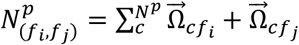. Aggregating all patients, we use the 25^th^ percentile as a cutoff value and remove all patients with 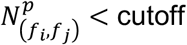. We find that removing these patients also removes the patients with a relatively high standard error.

#### Step 4: Identify significant CP-pairs (and microdomains) by comparing PMI distributions between recurring and non-recurring cohorts

For each CP-pair, (*f_i_, f_j_*), we compute the Kendall rank correlation, *τ*, (and p-value, *p_τ_*) between PMI(*f_i_, f_j_*) and time to recurrence (reported in days). In addition, we perform a Wilcoxon ranksum (WRS) test (does not reject the null hypothesis, *h_WRS_* = 0, or does reject the null hypothesis, *h_WRS_* = 1 and corresponding p-value *p_WRS_*) to determine if the NED-8yrs and REC-3yrs PMI(*f_i_, f_j_*) values are from equivalent distributions. We define a phenotype pair as significant if |*τ*| > *prctile*(*τ*, 99*th*), *p_τ_* ≤ 0.05, *h_WRS_* = 1, and *p_WRS_* ≤ 0.05. Each CP-pair distribution is shown in Figure S9. The Kendall rank correlation and Wilcoxon rank-sum test results are reported for CP-pair (and clinical variables) in Table S4. The 5 CP-pairs identified as significant by our analysis form two microdomains: 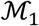: CP1-CP1, 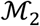: CP13-CP9-CP7-CP6-CP5.

#### Step 5: Perform partial correlation analysis to construct outcome-specific biomarker networks within each microdomain

For each microdomain, we aim to identify a biomarker network representation for the NED-8yrs and REC-3yrs patient groups. Within a single microdomain and for each patient cohort, we operate on cells with specialized states for the simplicity of network derivation (although ownership probabilities can be incorporated to include transitional and multi-transitional cell states into the network derivation). We then compute the partial correlation between each biomarker-pair, controlling for the remaining confounding biomarkers [1]. First, from the biomarker data (*X*: *N* × *p*) composed of *N* cells in the specific microdomain and *p* biomarkers, we compute the Pearson’s correlation coefficient matrix, *C*: *p* × *p*. Second, we invert the correlation coefficient matrix to obtain the precision matrix, Φ = *C*^−1^. Third, we compute the partial correlation coefficient between each biomarker pair, *ρ_ij_*, through normalization: 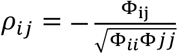. The partial correlation coefficients range from −1 to 1 representing the true correlation between each biomarker pair when all other confounding factors (other biomarkers) are removed [37]. For the sake of simplicity, we chose to pool cells across the two patient outcome cohorts: total number of cells in NED-8yrs cohort = 89,963 and REC-3yrs cohort = 89,596 (Table S3). However, one could also construct patient-specific networks, and based on the number of cells an additional step of matrix regularization may have to be incorporated to avoid overfitting.

To compare the partial correlation networks between the cohorts, NED-8yrs (*ρ^NED^*) and REC-3yrs (*ρ^REC^*), we use differential connectivity analysis with a permutation test [38]. In particular, we define a symmetric differential connectivity matrix, Δ_0_: *p* × *p*, where each entry, (*i, j*), represents the differential connectivity between a biomarker pair: 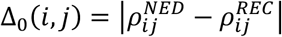. To test the significance of the differential, we use a permutation test (with *B* = 10,000 permutations). For one permutation iteration, the cells from both cohorts are aggregated and randomly sampled to form two data sets, mirroring the same number of cells as the NED-8yrs and REC-3yrs cohorts. Each permutation, *k*, results in a differential connectivity matrix of the permuted data, Δ_*k*_. The significance value (p-value) for a specific biomarker-pair (*i, j*) is computed as 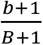 where 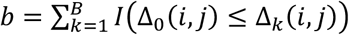. We define the biomarker-pairs with a differential connectivity score in the top 99^th^ percentile and p-value below 0.001 as significant (Table S5, S6).

## Supplementary Materials

### Supplementary Table Captions

**Table S1 –.** Previous work. We outline the state-of-the-art cellular phenotyping methods and analysis platforms/studies within the past decade specifying the data investigated, purpose of the study, and integration of the following components: spatial context, soft clustering, hierarchical component, and unsupervised techniques. The data generation methods include the following high-dimensional techniques: mass cytometry (CyTOF) [1], single-cell RNA sequencing (scRNA-seq) [2], multi to hyper – plexed immunofluorescence imaging (MxIF/HxIF) [3, 4], imaging mass cytometry (IMC) [5], small molecule fluorescent in situ hybridization (smFISH) [6], and flow cytometry [7]. Many single-cell clustering algorithms are supervised methods subjected to user-defined number of clusters and/or approximated ground truth reliant on prior biological knowledge [8–11]. Popular methods, PhenoGraph [12] and SPADE [13], dissect cell clusters from CyTOF data but lack a spatial component. All the analysis methods listed assume each cell belongs to exactly one celltype (hard-clustering) leaving no room to identify cell states along a cellular phenotypic continuum (transitional and multi-transitional cell states). To our knowledge, there does not exist a cellular phenotyping method with a combined spatial analysis framework which is unsupervised, harnesses a spatial component, and promotes the identification of the cellular phenotypic continuum.

| Previous method | Year | Input data | Purpose | Spatial context | Soft clustering | Hierarchical component | Unsupervised |
| --- | --- | --- | --- | --- | --- | --- | --- |
| Jackson, H.W., et al. [15] | 2020 | IMC | Spatially resolved, single-cell analysis to characterize intratumor phenotypic heterogeneity | Yes | No | No | No |
| Spatial-LDA [9] | 2020 | MxIF/HxIF | Recover microenvironment signatures | Yes | No | No | No |
| PhMED [8] | 2020 | CytoF, scRNA-seq | Embedding a 'manifold of manifolds' | No | No | Yes | No |
| BAMM-SC [16] | 2019 | scRNA-seq | Cluster scRNA-seq data from multiple individuals simultaneously | No | No | No | Yes |
| Zhu, Q., et al. [10] | 2018 | scRNAseq, smFISH | Distinguish between intrinsic and extrinsic effects of global gene expression | Yes | No | No | No |
| McKinley, E.T., et al. [17] | 2017 | HxIF | Quantify tuft cell number and distribution throughout the mouse small intestine and colon | Yes | No | No | No |
| Santamaria-Pang, A., et al. [11] | 2017 | HxIF | Machine-learning based method for classifying immune cell types in human tissue | No | No | Yes | No |
| X-shift [18] | 2016 | CytoF | Automated cell-subset clustering | No | No | No | Yes |
| Cytofkit [19] | 2016 | CytoF | Platform integrating bioinformatics methods and in-house novel algorithms for cytometry data analysis | No | No | No | No |
| PhenoGraph [12] | 2015 | CytoF | Algorithmically defines phenotypes in high-dimensional single cell data | No | No | No | Yes |
| viSNE [20] | 2013 | CytoF | Tool to map high-dimensional data onto two dimensions | No | No | No | No |
| SPADE [13] | 2011 | Flow cytometry, CytoF | Extract a hierarchy of cellular heterogeneity | No | No | Yes | Yes |

**Table S2 –.**
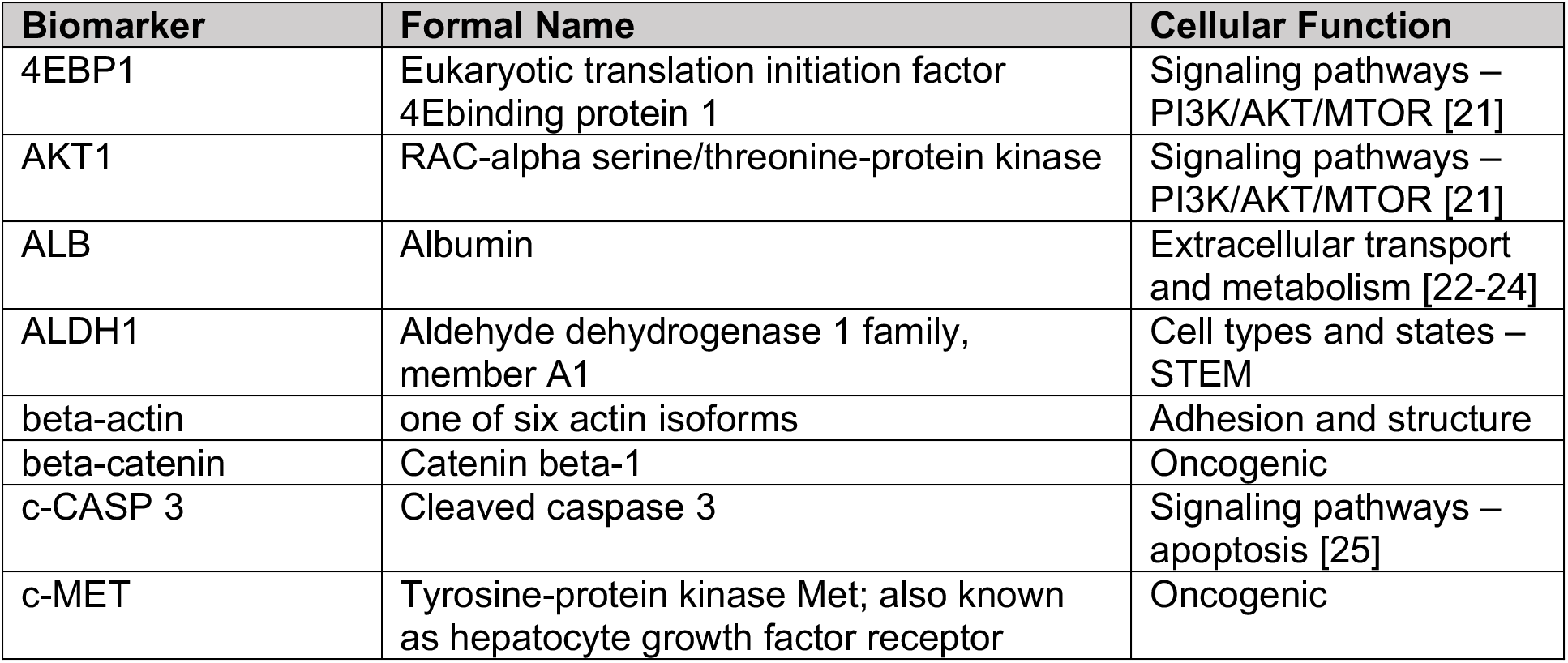
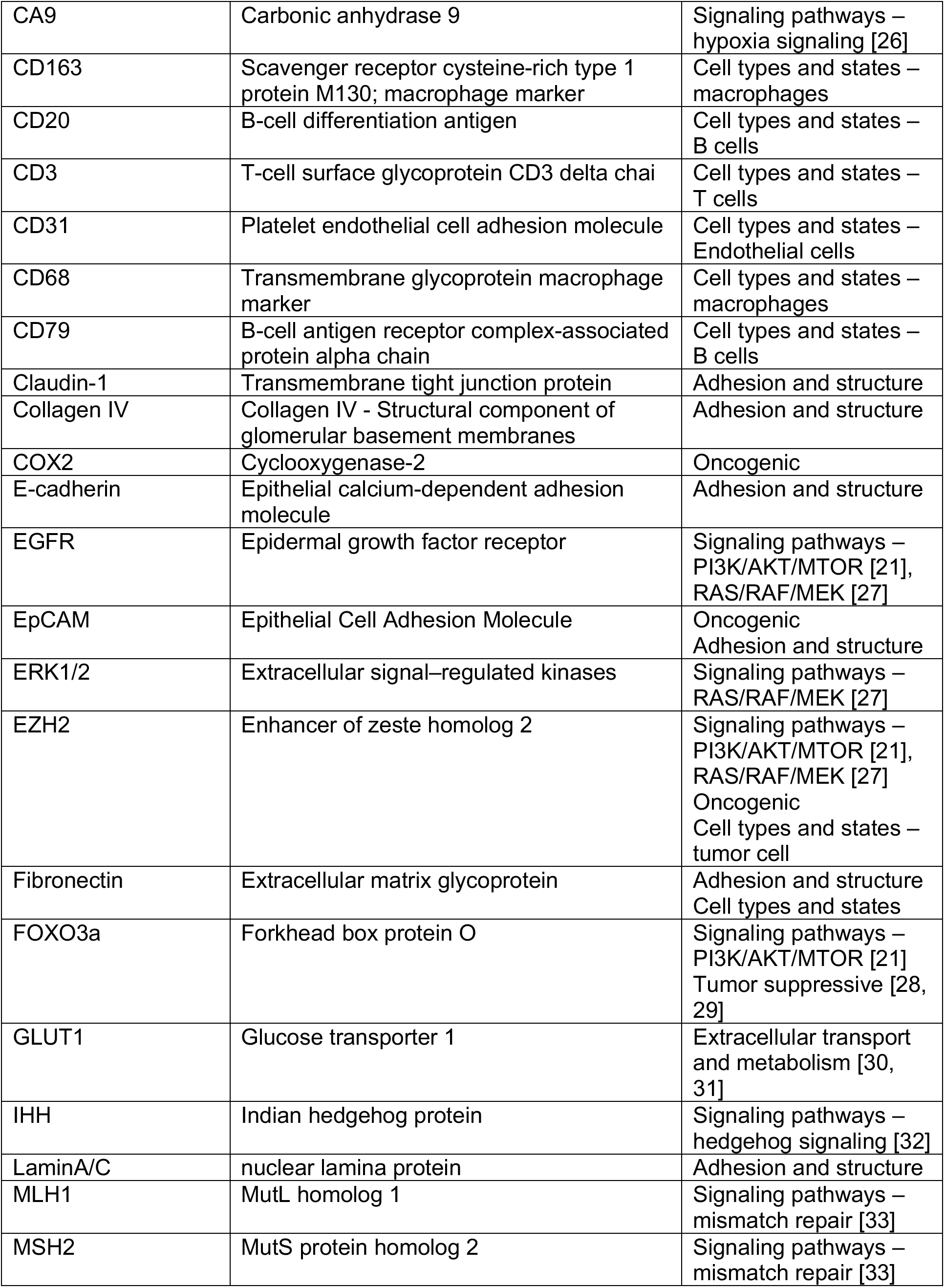
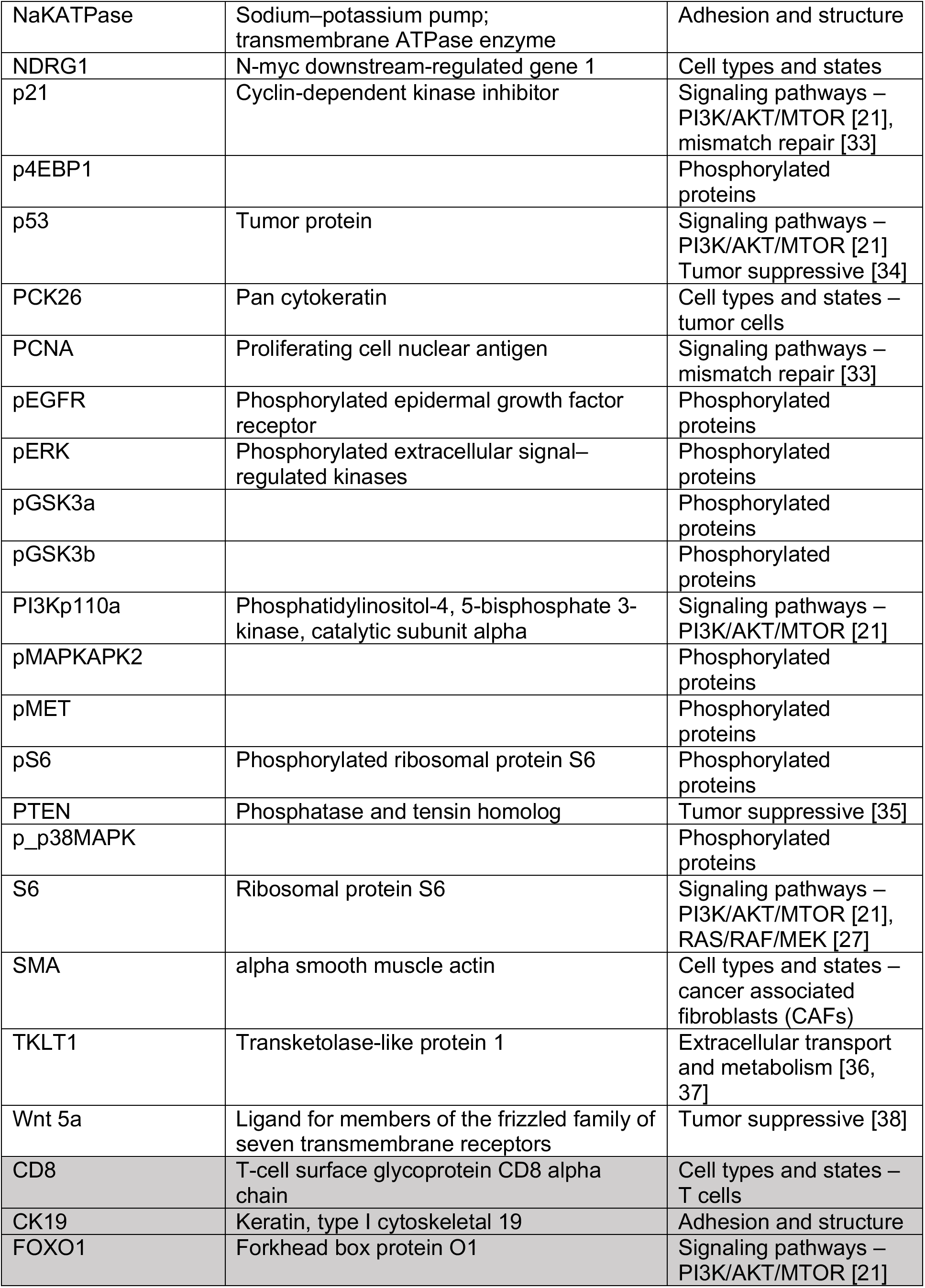
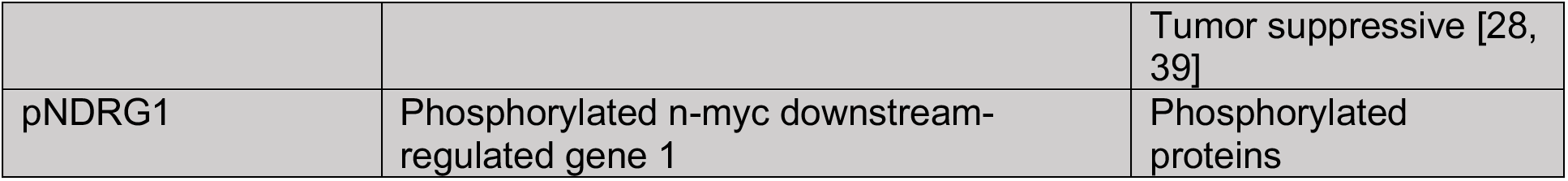
CRC biomarkers. List of the biomarkers in the CRC hyperplexed data set. Grayed biomarkers are eliminated due to quality control.

**Table S3 –.** CRC patient statistics. Patient statistics for the clinical data of the selected tissue samples.

| DATA | All data | NED-5yrs | REC-5yrs | NED-8yrs | REC-3yrs |
| --- | --- | --- | --- | --- | --- |
| # patients (%) | 213 (100%) | 154 (72.3%) | 59 (27.7%) | 45 (21.1%) | 46 (21.6%) |
| # males (%) | 105 (49.3%) | 73 (34.3%) | 32 (15%) | 15 (7%) | 28 (13.1%) |
| Mean age (std) | 63.6 (7.2) | 63 (7.1) | 65.4 (7.2) | 62.8 (6.4) | 65.4 (7.2) |
| Mean time-to-recurrence (std) | 1922.3 (1142.1) | 2393.3 (951.2) | 692.7 (504.4) | 3582.8 (469.3) | 464.6 (278.1) |
| # Stage I (%) | 94 (44.1%) | 91 (42.7%) | 3 (1.4%) | 25 (11.7%) | 2 (0.9%) |
| # Stage II (%) | 75 (35.2%) | 55 (25.8%) | 20 (9.4%) | 18 (8.5%) | 15 (7%) |
| # Stage III (%) | 44 (20.7%) | 8 (3.8%) | 36 (16.9%) | 2 (0.9%) | 29 (13.6%) |
| Mean # cells per tissue sample (std) | 2082.7 (513.4) | 2103 (492.3) | 2029.9 (565.7) | 1999.2 (476.8) | 1947.7 (593.9) |
| Total # of cells | 443618 | 323856 | 119762 | 89963 | 89596 |

**Table S4 –.** Correlation of pointwise mutual information with time-to-recurrence. Correlation analysis with the clinical information (sex, grade, stage, age), CP fractions per tissue sample, and CP-pair pointwise mutual information (PMI) values. We compute the Kendall rank correlation coefficient, *τ*, (and significance, *p_τ_*) between each covariate and the time-to-recurrence (days). In addition, we perform the Wilcoxon rank-sum test (hypothesis *h_WRS_* and significance, *p_WRS_*) to determine if for each covariate, the NED-8yrs and REC-3yrs patient tissue samples originate from the same underlying distribution. Covariates are considered significant (highlighted in yellow) if *h_WRS_* = 1 (*p_WRS_* ≤ 0.05) and |*τ*| > 0.1 (*p_τ_* ≤ 0.05).

| Covariate | Correlation coefficient (Kendall rank, $\tau$) | Correlation coefficient significance ($p_\tau$) | Wilcoxon rank-sum test ($h_{WRS}$) | Wilcoxon rank-sum test significance ($p_{WRS}$) | # NED-8yrs | # REC-3yrs |
| --- | --- | --- | --- | --- | --- | --- |
| stage | -0.449 | 0 | 0 | 1 | 45 | 46 |
| grade | -0.377 | 0 | 0 | 1 | 45 | 45 |
| CP13::CP5 | -0.251 | 0.0029 | 0.0362 | 1 | 35 | 31 |
| CP9::CP6 | -0.216 | 0.0161 | 0.0302 | 1 | 32 | 27 |
| CP9::CP5 | -0.215 | 0.0051 | 0.005 | 1 | 40 | 39 |
| sex | 0.207 | 0.0169 | 0.009 | 1 | 45 | 46 |
| CP1::CP1 | -0.188 | 0.0376 | 0.014 | 1 | 29 | 29 |
| CP13::CP6 | -0.183 | 0.0572 | 0.0446 | 1 | 29 | 23 |
| CP9::CP7 | -0.159 | 0.0259 | 0.0198 | 1 | 45 | 46 |
| CP5::CP1 | -0.148 | 0.0668 | 0.0211 | 1 | 34 | 38 |
| CP7::CP6 | -0.14 | 0.0903 | 0.0426 | 1 | 37 | 32 |
| CP8::CP4 | 0.139 | 0.1181 | 0.0475 | 1 | 31 | 29 |
| CP12::CP6 | -0.124 | 0.1512 | 0.0337 | 1 | 32 | 31 |
| CP4::CP2 | 0.053 | 0.5042 | 0.0236 | 1 | 36 | 39 |
| CP12::CP5 | -0.245 | 0.0018 | 0.069 | 0 | 39 | 37 |
| CP12::CP3 | -0.205 | 0.011 | 0.0519 | 0 | 39 | 33 |
| CP13::CP3 | -0.188 | 0.0219 | 0.0703 | 0 | 38 | 32 |
| CP5::CP5 | -0.161 | 0.1079 | 0.1349 | 0 | 28 | 20 |
| CP13::CP13 | 0.159 | 0.0441 | 0.0556 | 0 | 39 | 36 |
| CP7::CP5 | -0.15 | 0.0515 | 0.1481 | 0 | 39 | 40 |
| age | -0.148 | 0.0427 | 0.0558 | 0 | 45 | 46 |
| CP9::CP3 | -0.147 | 0.0727 | 0.315 | 0 | 36 | 34 |
| CP5::CP2 | -0.146 | 0.0588 | 0.2096 | 0 | 40 | 38 |
| CP10::CP5 | -0.146 | 0.0814 | 0.0626 | 0 | 34 | 33 |
| CP13::CP8 | 0.144 | 0.0486 | 0.489 | 0 | 43 | 44 |
| CP7::CP3 | -0.137 | 0.0873 | 0.2389 | 0 | 39 | 34 |
| CP8 | 0.133 | 0.0632 | 0.3979 | 0 | 45 | 46 |
| CP5 | -0.132 | 0.0656 | 0.248 | 0 | 45 | 46 |
| CP6::CP2 | -0.127 | 0.1208 | 0.1183 | 0 | 38 | 32 |
| CP11::CP5 | -0.125 | 0.1283 | 0.1485 | 0 | 35 | 35 |
| CP8::CP5 | -0.124 | 0.1512 | 0.0973 | 0 | 33 | 30 |
| CP13 | 0.117 | 0.1013 | 0.2547 | 0 | 45 | 46 |
| CP13::CP11 | 0.115 | 0.112 | 0.3541 | 0 | 45 | 43 |
| CP11::CP4 | 0.112 | 0.1924 | 0.3034 | 0 | 34 | 30 |
| CP11::CP3 | -0.111 | 0.1798 | 0.5444 | 0 | 35 | 34 |
| CP9::CP9 | -0.11 | 0.1719 | 0.2111 | 0 | 37 | 35 |
| CP8::CP7 | 0.109 | 0.1377 | 0.3872 | 0 | 41 | 45 |
| CP6::CP3 | 0.108 | 0.2506 | 0.2682 | 0 | 27 | 27 |
| CP6::CP5 | -0.106 | 0.2431 | 0.0836 | 0 | 31 | 27 |
| CP3::CP3 | -0.103 | 0.2761 | 0.2181 | 0 | 25 | 29 |
| CP11::CP9 | -0.101 | 0.1616 | 0.0512 | 0 | 43 | 46 |
| CP5::CP3 | -0.099 | 0.2458 | 0.4745 | 0 | 33 | 32 |
| CP9 | -0.096 | 0.179 | 0.0857 | 0 | 45 | 46 |
| CP2::CP1 | -0.095 | 0.1919 | 0.2679 | 0 | 43 | 44 |
| CP8::CP3 | -0.094 | 0.2873 | 0.4483 | 0 | 32 | 29 |
| CP10::CP6 | -0.094 | 0.3016 | 0.1345 | 0 | 31 | 27 |
| CP10::CP8 | 0.093 | 0.2235 | 0.9231 | 0 | 37 | 43 |
| CP10::CP3 | -0.092 | 0.2838 | 0.7524 | 0 | 32 | 32 |
| CP11::CP7 | -0.09 | 0.2135 | 0.3187 | 0 | 44 | 45 |
| CP11::CP10 | 0.089 | 0.224 | 0.554 | 0 | 42 | 44 |
| CP12::CP8 | 0.089 | 0.2188 | 0.3391 | 0 | 44 | 45 |
| CP10::CP4 | 0.084 | 0.3194 | 0.051 | 0 | 35 | 32 |
| CP13::CP12 | 0.081 | 0.2654 | 0.4602 | 0 | 44 | 44 |
| CP11::CP6 | -0.077 | 0.3818 | 0.2568 | 0 | 32 | 30 |
| CP3 | -0.071 | 0.3183 | 0.7298 | 0 | 45 | 46 |
| CP10::CP9 | -0.069 | 0.3404 | 0.1161 | 0 | 44 | 45 |
| CP3::CP2 | -0.069 | 0.3848 | 0.787 | 0 | 38 | 37 |
| CP8::CP6 | -0.069 | 0.5096 | 0.2337 | 0 | 25 | 21 |
| CP4 | 0.069 | 0.3387 | 0.0734 | 0 | 45 | 46 |
| CP13::CP4 | 0.065 | 0.4365 | 0.2406 | 0 | 36 | 32 |
| CP9::CP2 | -0.065 | 0.3685 | 0.3835 | 0 | 45 | 45 |
| CP12::CP7 | -0.063 | 0.3853 | 0.2842 | 0 | 45 | 44 |
| CP8::CP2 | 0.063 | 0.4019 | 0.1206 | 0 | 42 | 42 |
| CP7::CP1 | -0.062 | 0.3874 | 0.1232 | 0 | 45 | 45 |
| CP6::CP6 | -0.061 | 0.5999 | 0.4053 | 0 | 24 | 14 |
| CP13::CP2 | 0.061 | 0.401 | 0.2891 | 0 | 45 | 43 |
| CP4::CP1 | -0.059 | 0.4467 | 0.6284 | 0 | 39 | 38 |
| CP11::CP1 | -0.059 | 0.4276 | 0.0975 | 0 | 42 | 43 |
| CP8::CP1 | 0.057 | 0.4755 | 0.9789 | 0 | 37 | 38 |
| CP12::CP4 | 0.054 | 0.5099 | 0.4209 | 0 | 36 | 34 |
| CP7::CP4 | 0.054 | 0.4984 | 0.2092 | 0 | 38 | 37 |
| CP10 | 0.054 | 0.4507 | 0.9588 | 0 | 45 | 46 |
| CP7::CP2 | -0.053 | 0.4632 | 0.8793 | 0 | 45 | 44 |
| CP8::CP8 | 0.052 | 0.5617 | 0.5247 | 0 | 29 | 31 |
| CP11::CP8 | 0.051 | 0.4938 | 0.7325 | 0 | 40 | 43 |
| CP13::CP10 | 0.05 | 0.5039 | 0.7678 | 0 | 41 | 43 |
| CP9::CP1 | -0.05 | 0.5088 | 0.1943 | 0 | 42 | 41 |
| CP1 | -0.05 | 0.4886 | 0.0871 | 0 | 45 | 46 |
| CP12::CP9 | -0.049 | 0.499 | 0.4338 | 0 | 45 | 45 |
| CP6 | -0.048 | 0.4993 | 0.821 | 0 | 45 | 46 |
| CP3::CP1 | -0.047 | 0.58 | 0.4152 | 0 | 32 | 34 |
| CP6::CP1 | -0.047 | 0.6224 | 0.7488 | 0 | 28 | 26 |
| CP7 | -0.045 | 0.5282 | 0.921 | 0 | 45 | 46 |
| CP4::CP4 | 0.045 | 0.6515 | 0.2989 | 0 | 24 | 26 |
| CP12::CP1 | -0.041 | 0.5809 | 0.1746 | 0 | 41 | 45 |
| CP11 | 0.041 | 0.5693 | 0.9273 | 0 | 45 | 46 |
| CP9::CP8 | 0.04 | 0.5813 | 0.6048 | 0 | 43 | 45 |
| CP10::CP10 | 0.037 | 0.6441 | 0.6156 | 0 | 37 | 36 |
| CP6::CP4 | -0.035 | 0.6885 | 0.3542 | 0 | 34 | 28 |
| CP7::CP7 | -0.035 | 0.6569 | 0.9148 | 0 | 39 | 38 |
| CP5::CP4 | -0.034 | 0.6961 | 0.6987 | 0 | 33 | 32 |
| CP9::CP4 | -0.031 | 0.7008 | 0.795 | 0 | 39 | 36 |
| CP13::CP7 | 0.03 | 0.6758 | 0.6283 | 0 | 45 | 45 |
| CP11::CP2 | -0.03 | 0.6838 | 0.6642 | 0 | 45 | 43 |
| CP11::CP11 | 0.027 | 0.7353 | 0.555 | 0 | 36 | 37 |
| CP2 | 0.027 | 0.7112 | 0.2716 | 0 | 45 | 46 |
| CP12::CP12 | -0.026 | 0.743 | 0.8651 | 0 | 39 | 39 |
| CP12::CP10 | 0.025 | 0.7304 | 0.9155 | 0 | 44 | 43 |
| CP4::CP3 | 0.025 | 0.7735 | 0.6628 | 0 | 33 | 33 |
| CP12::CP11 | -0.021 | 0.7687 | 0.86 | 0 | 44 | 45 |
| CP2::CP2 | 0.021 | 0.8035 | 0.8825 | 0 | 31 | 37 |
| CP12 | -0.016 | 0.8263 | 0.8707 | 0 | 45 | 46 |
| CP13::CP1 | -0.013 | 0.862 | 0.4338 | 0 | 42 | 42 |
| CP10::CP2 | 0.01 | 0.8882 | 0.701 | 0 | 45 | 43 |
| CP10::CP1 | -0.006 | 0.9382 | 0.2922 | 0 | 40 | 41 |
| CP10::CP7 | 0.005 | 0.9463 | 0.8535 | 0 | 45 | 44 |
| CP12::CP2 | -0.005 | 0.9474 | 0.6805 | 0 | 44 | 43 |
| CP13::CP9 | 0.003 | 0.9684 | 0.7291 | 0 | 44 | 44 |

**Table S5 –.** Microdomain 1 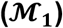 specific biomarker interaction networks with partial correlation analysis. Within each spatial microdomain (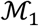: CP1::CP1, 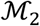: CP13::CP9::CP7::CP6::CP5) we compute the partial correlation network (1,275 pairs total) for the NED-8yrs and REC-3yrs patient groups and compute the differential (Δ_0_) between the cohorts with a corresponding p value using a permutation test (see Methods). The partial correlation values represent the true correlation between two biomarkers when all other confounding factors (other biomarkers) are removed. This table reports the biomarker pairs with a significant differential in the 99^th^ percentile within 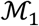.

| Biomarker-pair | NED-8yrs<br>partial<br>correlation | REC-3yrs<br>partial<br>correlation | $\Delta_0$ | p-value ($\Delta_0$) |
| --- | --- | --- | --- | --- |
| ALDH1::CD68 | 0.137 | -0.378 | 0.515 | 0.0001 |
| CD163::PI3Kp110a | -0.026 | 0.428 | 0.454 | 0.0001 |
| Wnt5a::IHH | 0.337 | -0.057 | 0.394 | 0.0001 |
| pMET::CD163 | 0.165 | -0.191 | 0.357 | 0.0001 |
| EZH2::ALDH1 | 0.163 | -0.192 | 0.355 | 0.0001 |
| CA9::PI3Kp110a | -0.044 | 0.308 | 0.353 | 0.0001 |
| CD20::pEGFR | 0.082 | 0.425 | 0.344 | 0.0001 |
| p4EBP1::CA9 | 0.135 | -0.195 | 0.33 | 0.0001 |
| beta-catenin::EGFR | -0.11 | 0.206 | 0.316 | 0.0001 |
| CA9::CD163 | 0.051 | -0.256 | 0.308 | 0.0001 |
| EZH2::c-MET | 0.089 | 0.395 | 0.306 | 0.0001 |
| beta-catenin::ALDH1 | -0.227 | 0.071 | 0.298 | 0.0001 |
| Wnt5a::4EBP1 | 0.137 | -0.155 | 0.293 | 0.0001 |

**Table S6 –.** Microdomain 2 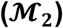 specific biomarker interaction networks with partial correlation analysis. With the same method as described in Table S5, we report the biomarker pairs with a significant differential in the 99^th^ percentile within microdomain 2 (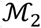: CP13::CP9::CP7::CP6::CP5).

| Biomarker-pair | NED-8yrs<br>partial<br>correlation | REC-3yrs<br>partial<br>correlation | $\Delta_0$ | p-value ($\Delta_0$) |
| --- | --- | --- | --- | --- |
| pERK::CD31 | 0.426 | 0.034 | 0.392 | 0.0001 |
| FOXO3a::PI3Kp110a | -0.209 | 0.111 | 0.32 | 0.0001 |
| Wnt5a::IHH | 0.203 | -0.098 | 0.301 | 0.0001 |
| MLH1::pGSK3a | 0.035 | 0.323 | 0.288 | 0.0001 |
| IHH::NDRG1 | -0.022 | 0.265 | 0.287 | 0.0001 |
| NDRG1::PCNA | 0.049 | -0.232 | 0.28 | 0.0001 |
| ALDH1::p53 | -0.347 | -0.067 | 0.28 | 0.0001 |
| p_p38MAPK::p53 | -0.112 | 0.159 | 0.271 | 0.0001 |
| p4EBP1::4EBP1 | 0.263 | -0.007 | 0.27 | 0.0001 |
| EGFR::4EBP1 | 0.006 | 0.274 | 0.267 | 0.0001 |
| ERK1/2::PCNA | -0.219 | 0.039 | 0.258 | 0.0001 |
| CD163::PI3Kp110a | 0.165 | 0.421 | 0.256 | 0.0001 |
| FOXO3a::4EBP1 | 0.041 | -0.213 | 0.255 | 0.0001 |

**Table S7 –.**
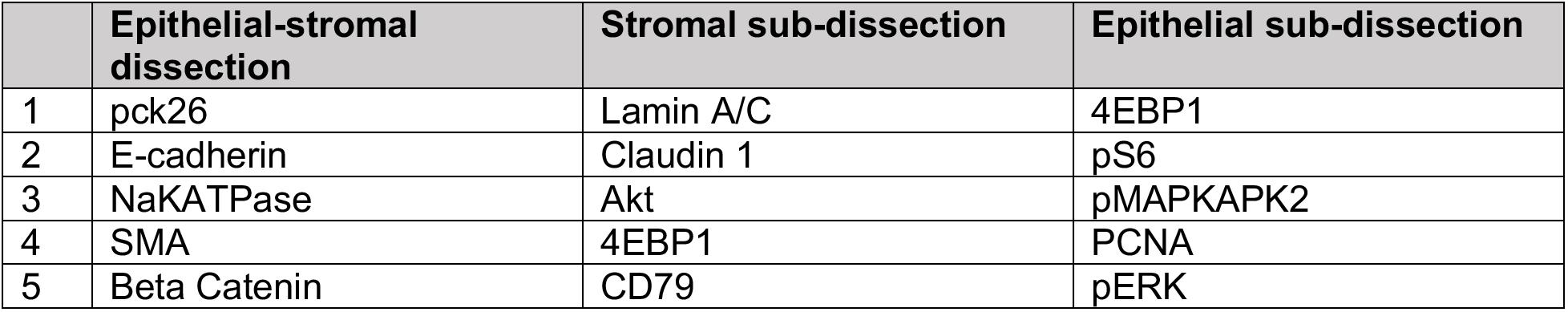
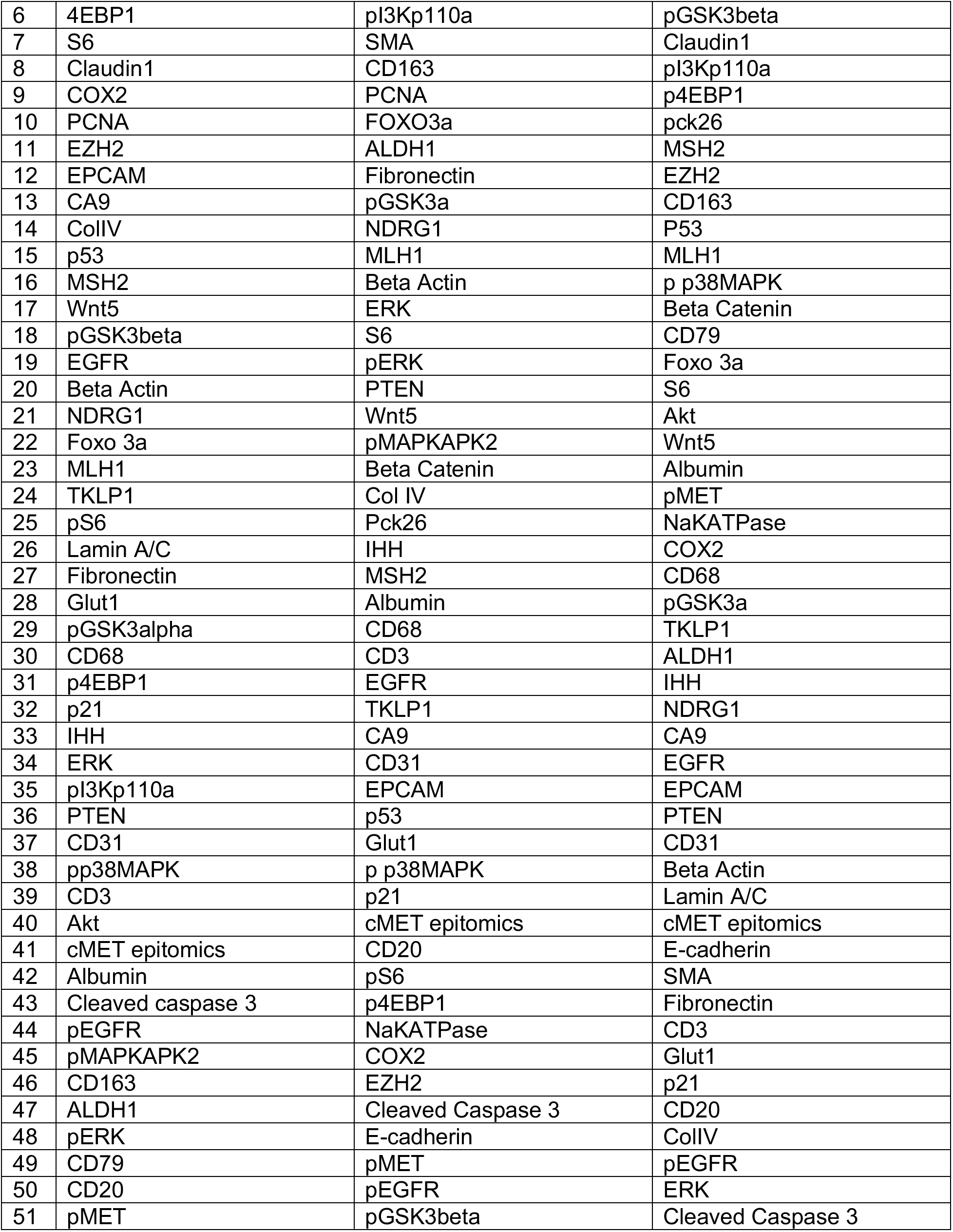
Biomarker order for virtual simulation. Biomarkers listed in order based on discriminative ranking for LEAPH performed on ALL-DATA (Figure 4). The epithelial-stromal dissection is from level 1 of the LEAPH hierarchy and the epithelial/stromal sub-dissection is from level 2. Each ranking is used to determine the order of adding biomarkers to LEAPH to reveal the minimal subset of biomarkers needed to reproduce the original computational phenotypes (Figure S10).

|  | Epithelial-stromal<br>dissection | Stromal sub-dissection | Epithelial sub-dissection |
| --- | --- | --- | --- |
| 1 | pck26 | Lamin A/C | 4EBP1 |
| 2 | E-cadherin | Claudin 1 | pS6 |
| 3 | NaKATPase | Akt | pMAPKAPK2 |
| 4 | SMA | 4EBP1 | PCNA |
| 5 | Beta Catenin | CD79 | pERK |
| 6 | 4EBP1 | pl3Kp110a | pGSK3beta |
| 7 | S6 | SMA | Claudin1 |
| 8 | Claudin1 | CD163 | pl3Kp110a |
| 9 | COX2 | PCNA | p4EBP1 |
| 10 | PCNA | FOXO3a | pck26 |
| 11 | EZH2 | ALDH1 | MSH2 |
| 12 | EPCAM | Fibronectin | EZH2 |
| 13 | CA9 | pGSK3a | CD163 |
| 14 | ColIV | NDRG1 | P53 |
| 15 | p53 | MLH1 | MLH1 |
| 16 | MSH2 | Beta Actin | p p38MAPK |
| 17 | Wnt5 | ERK | Beta Catenin |
| 18 | pGSK3beta | S6 | CD79 |
| 19 | EGFR | pERK | Foxo 3a |
| 20 | Beta Actin | PTEN | S6 |
| 21 | NDRG1 | Wnt5 | Akt |
| 22 | Foxo 3a | pMAPKAPK2 | Wnt5 |
| 23 | MLH1 | Beta Catenin | Albumin |
| 24 | TKLP1 | Col IV | pMET |
| 25 | pS6 | Pck26 | NaKATPase |
| 26 | Lamin A/C | IHH | COX2 |
| 27 | Fibronectin | MSH2 | CD68 |
| 28 | Glut1 | Albumin | pGSK3a |
| 29 | pGSK3alpha | CD68 | TKLP1 |
| 30 | CD68 | CD3 | ALDH1 |
| 31 | p4EBP1 | EGFR | IHH |
| 32 | p21 | TKLP1 | NDRG1 |
| 33 | IHH | CA9 | CA9 |
| 34 | ERK | CD31 | EGFR |
| 35 | pl3Kp110a | EPCAM | EPCAM |
| 36 | PTEN | p53 | PTEN |
| 37 | CD31 | Glut1 | CD31 |
| 38 | pp38MAPK | p p38MAPK | Beta Actin |
| 39 | CD3 | p21 | Lamin A/C |
| 40 | Akt | cMET epitomics | cMET epitomics |
| 41 | cMET epitomics | CD20 | E-cadherin |
| 42 | Albumin | pS6 | SMA |
| 43 | Cleaved caspase 3 | p4EBP1 | Fibronectin |
| 44 | pEGFR | NaKATPase | CD3 |
| 45 | pMAPKAPK2 | COX2 | Glut1 |
| 46 | CD163 | EZH2 | p21 |
| 47 | ALDH1 | Cleaved Caspase 3 | CD20 |
| 48 | pERK | E-cadherin | ColIV |
| 49 | CD79 | pMET | pEGFR |
| 50 | CD20 | pEGFR | ERK |
| 51 | pMET | pGSK3beta | Cleaved Caspase 3 |

### Supplementary Figure Captions

**Figure S1 –.**
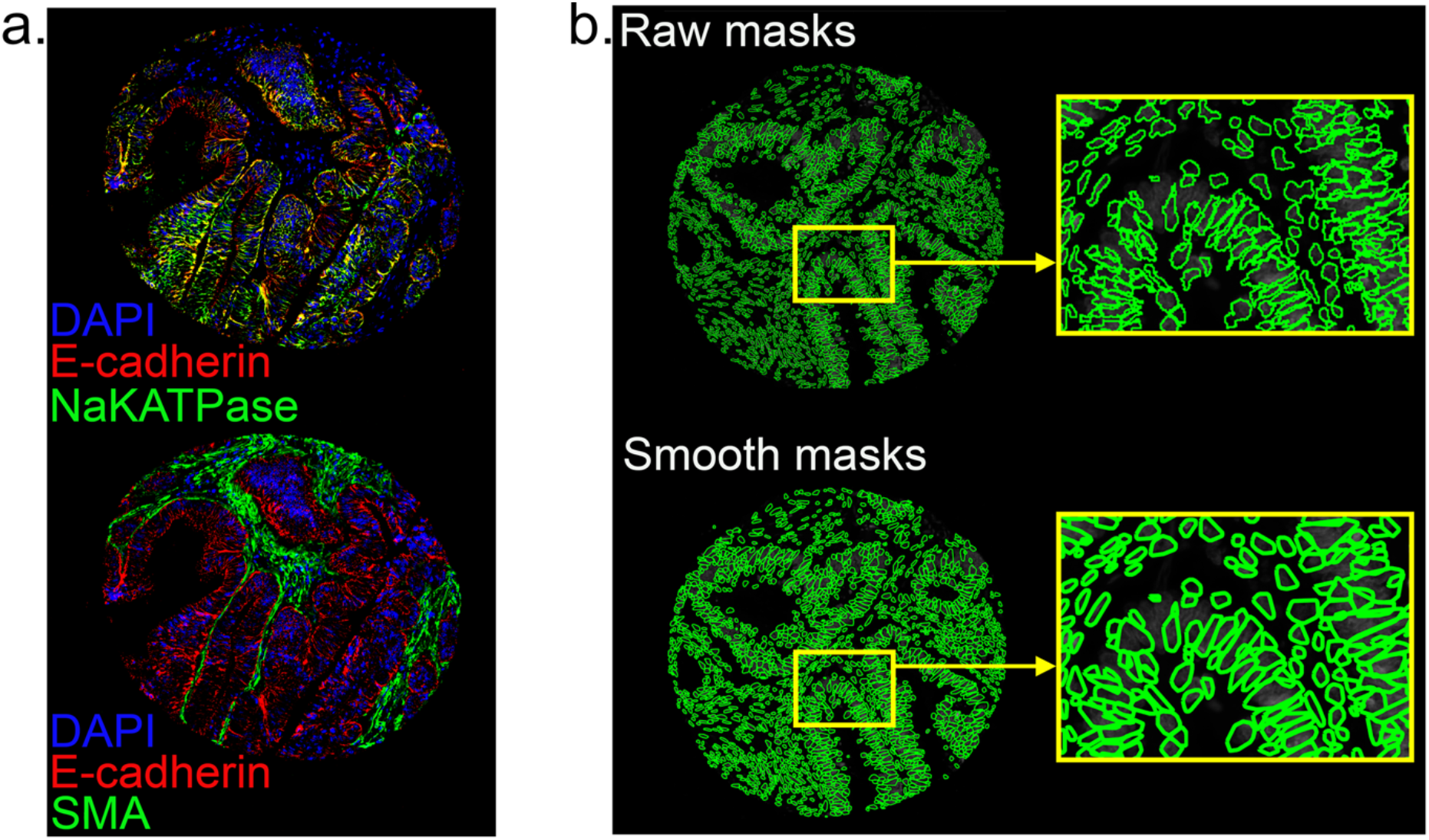
Cell segmentation. **a.** Pseudo colored image of a tissue sample with DAPI (blue), E-cadherin (red), NaKATPase (green, ***top***), and SMA (green, ***bottom***) biomarkers used for cell segmentation. **b**. The raw cell segmentation masks (***top***) and the smoothed cell masks used for visualization (***bottom***). The zoomed in regions of the raw and smoothed cell masks show a closer visual of the cell borders.

**Figure S2 –.**
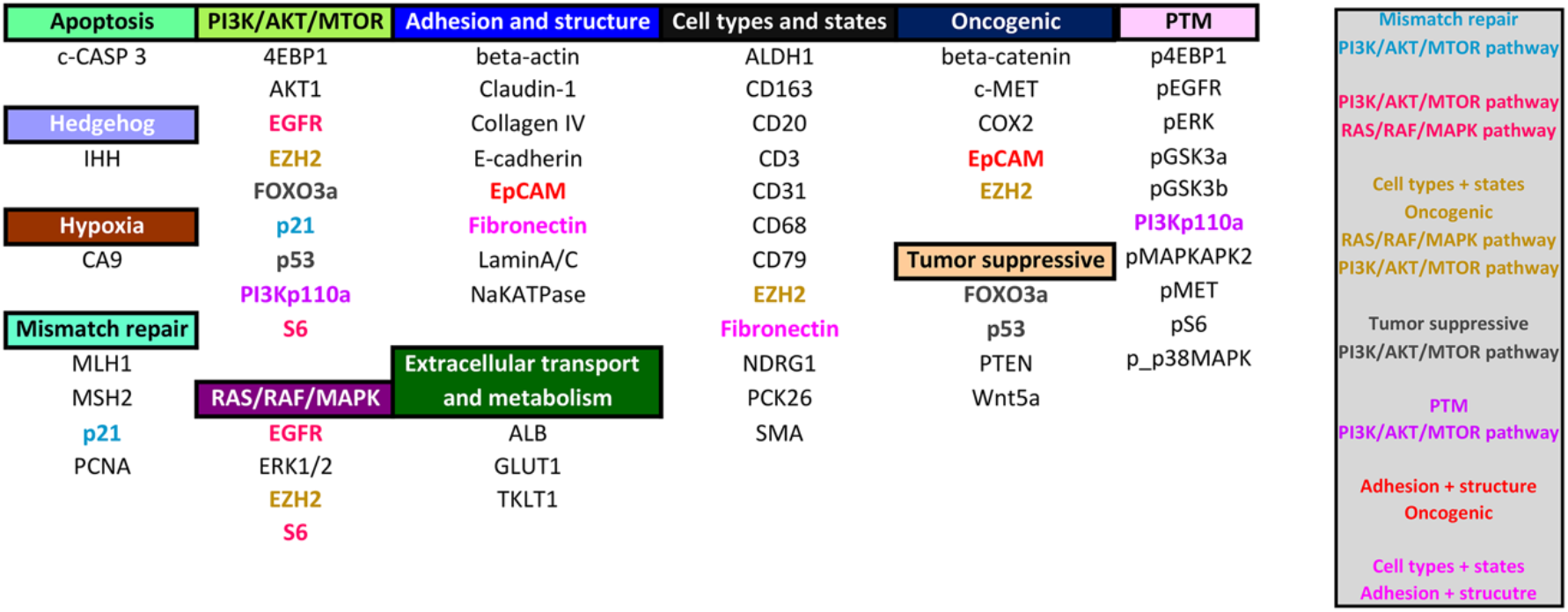
Biomarker functions and properties. List of the biomarkers in the CRC hyperplexed data set organized by their assumed cellular functions, or involvement in cellular processes, derived from the literature (see Table S1 for more information). Bolded biomarkers hold more than one cellular function (see legend in gray box).

**Figure S3 –.**
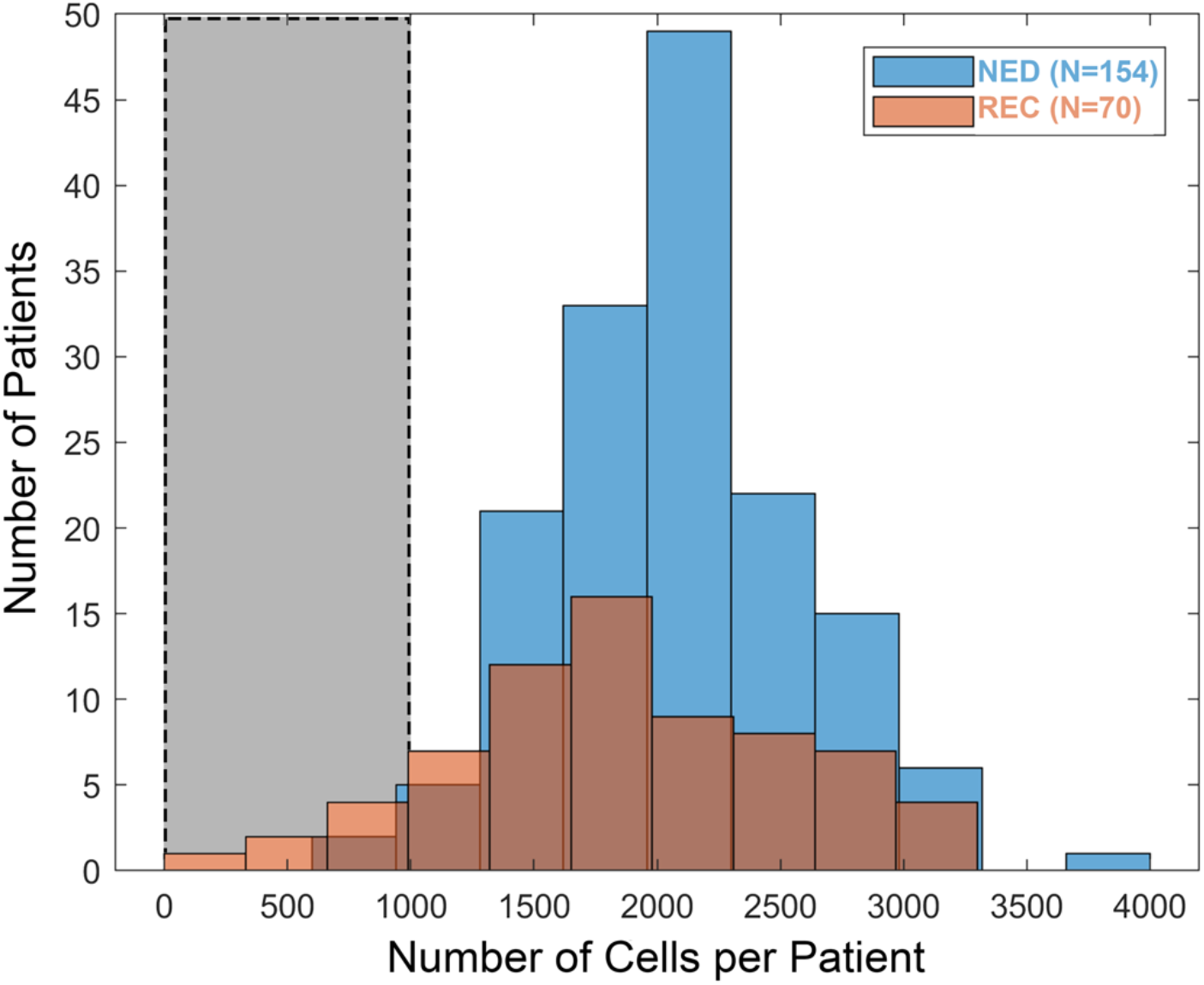
Cell count quality control. Tissue damage and other imaging artifacts impact biomarker staining and subsequent cell segmentation. We observe this variability in the number of cells segmented from each spot, shown here as a histogram. We remove all spots with less than a set 1000 cell threshold based on the 20^th^ percentile of the distribution (gray box).

**Figure S4 –.**
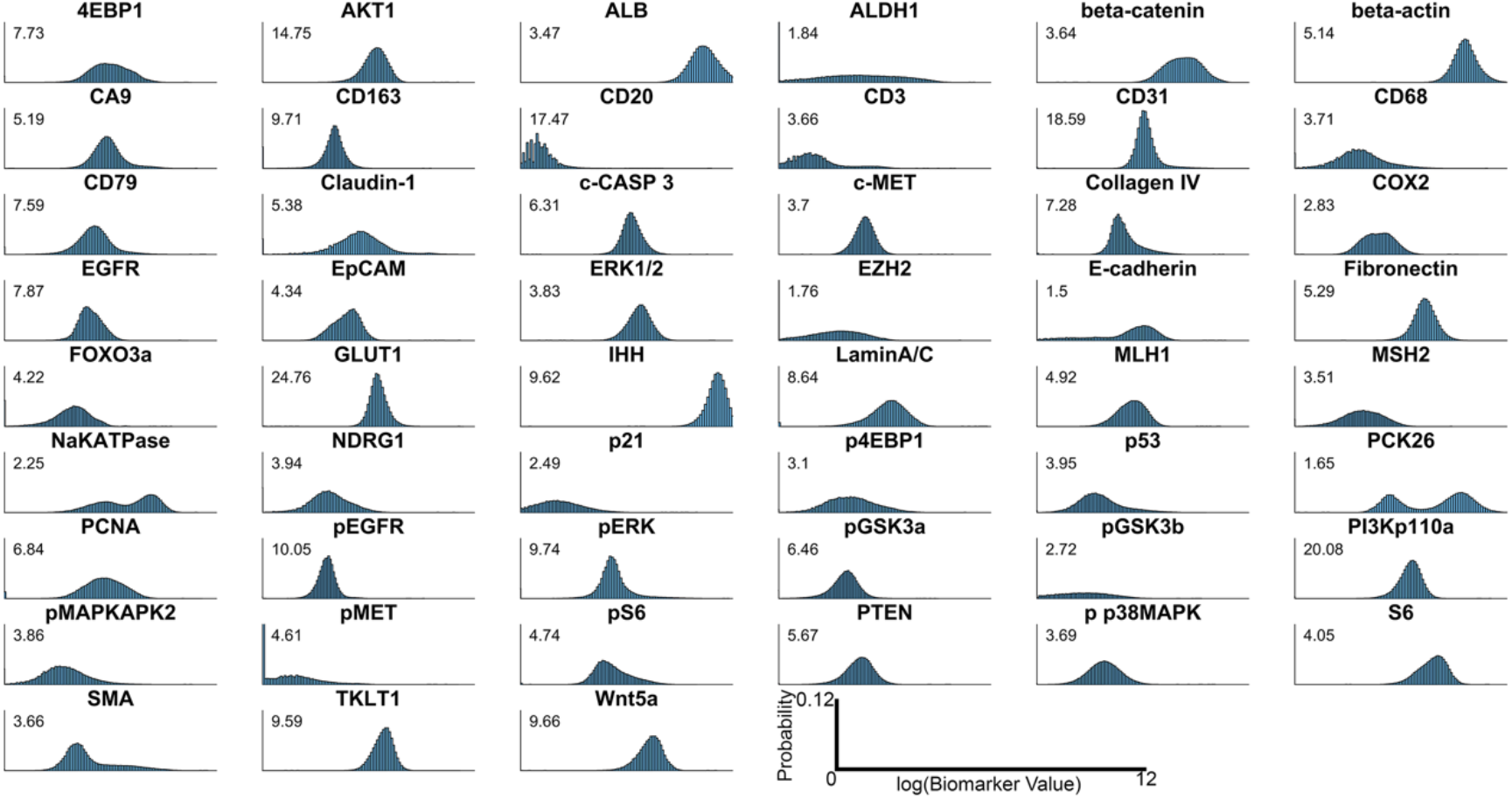
Non-normal distribution of log transformed biomarker values. Histograms of biomarker values over ~500K cells from 213 tissue samples. The x-axis is the biomarker values log transformed which range between 0 and 12. Each biomarker has a dynamic range of values, e.g., log(ALDH1):0-12 but log(CD20):0-4. The probability range (y-axis) is same across all plots (0-0.12). The values alongside each biomarker distribution represent the kurtosis. When compared to a Gaussian distribution (kurtosis=3), many of the biomarker distributions in our data are non-Gaussian with kurtosis values above (superGaussian) and below 3 (sub-Gaussian).

**Figure S5 –.**
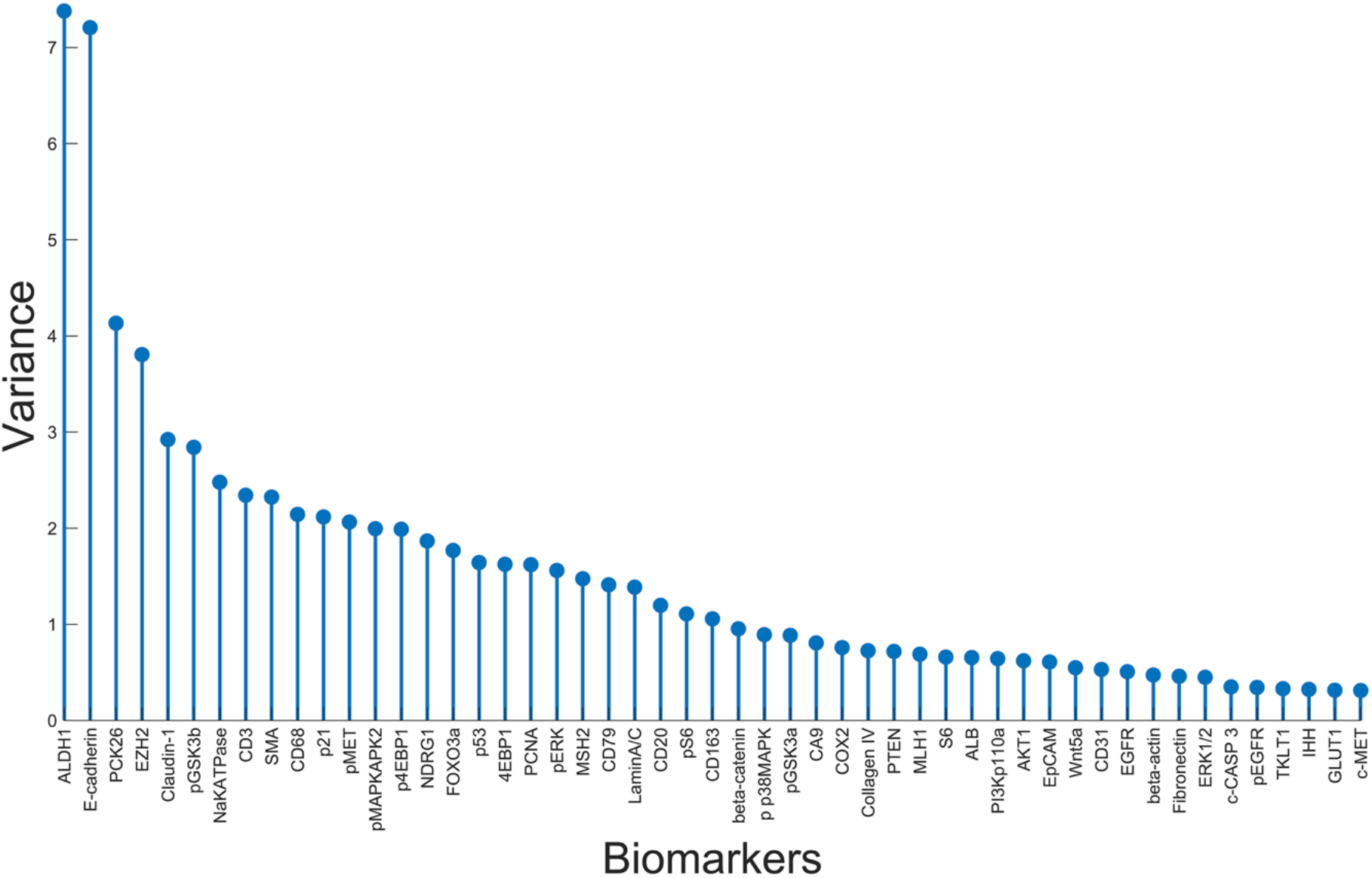
Variance for each biomarker distribution. Variance for each biomarker over the entire CRC data. We chose a two-dimensional latent space for the MFA model, as we observed this is enough to capture the input variance.

**Figure S6 –.**
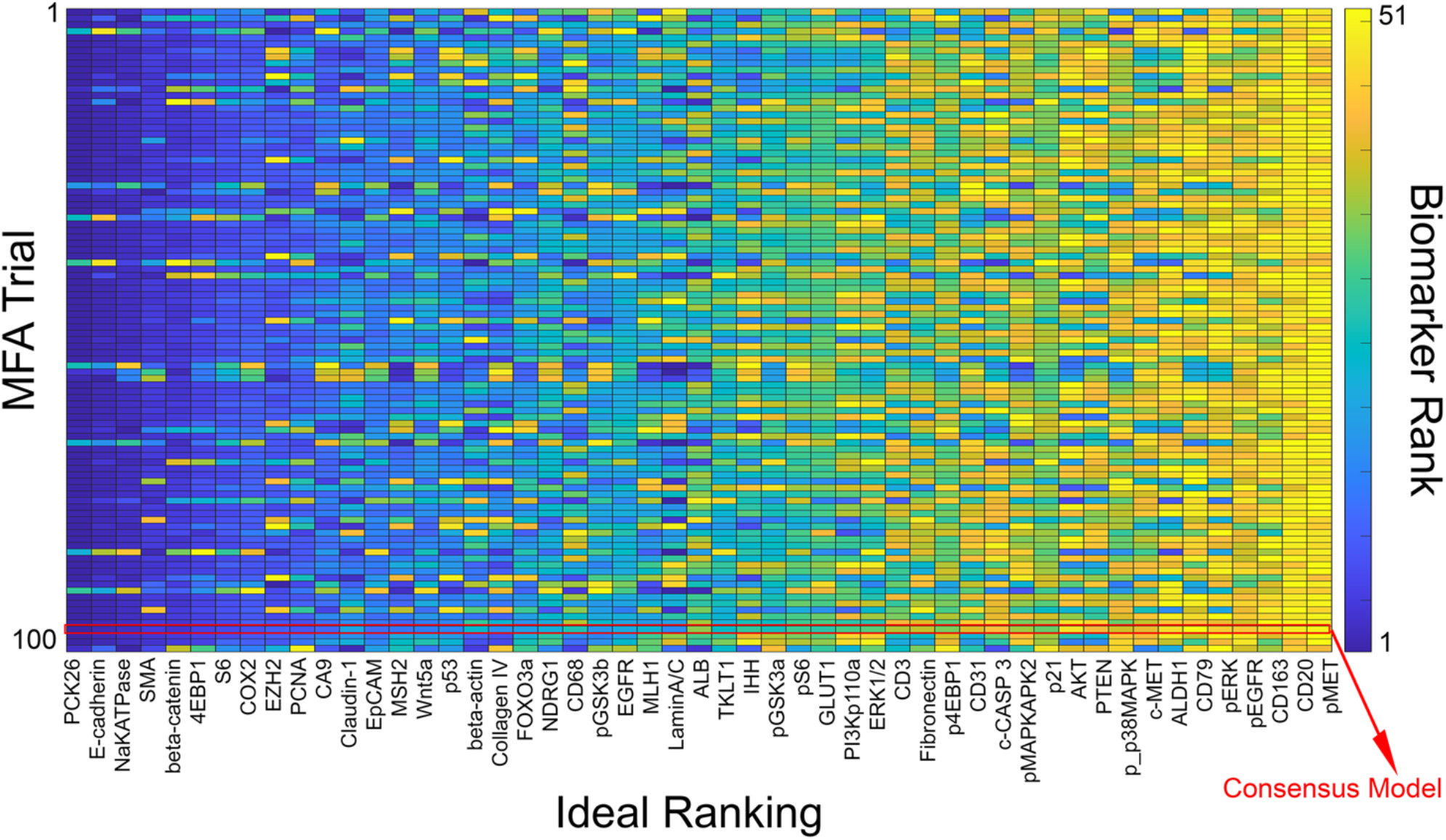
Mixture of Factor Analyzers (MFA) model selection. MFA model parameters are learned through the Expectation-Maximization (EM) algorithm with a random initialization and provides no guarantee to converge to a global minimum [14]. To account for this and ensure stability, we perform a hundred different EM optimizations, each initialized randomly. Each optimization yields an MFA model with a set of model parameters. We compute the biomarker ranking for each set of model parameters (see Methods) and aggregate all biomarker rankings to compute their mean ranking. The model with a biomarker ranking closest (Euclidean distance) to the mean ranking is selected as the consensus model and deemed to provide an optimal subspace representation.

**Figure S7 –.**
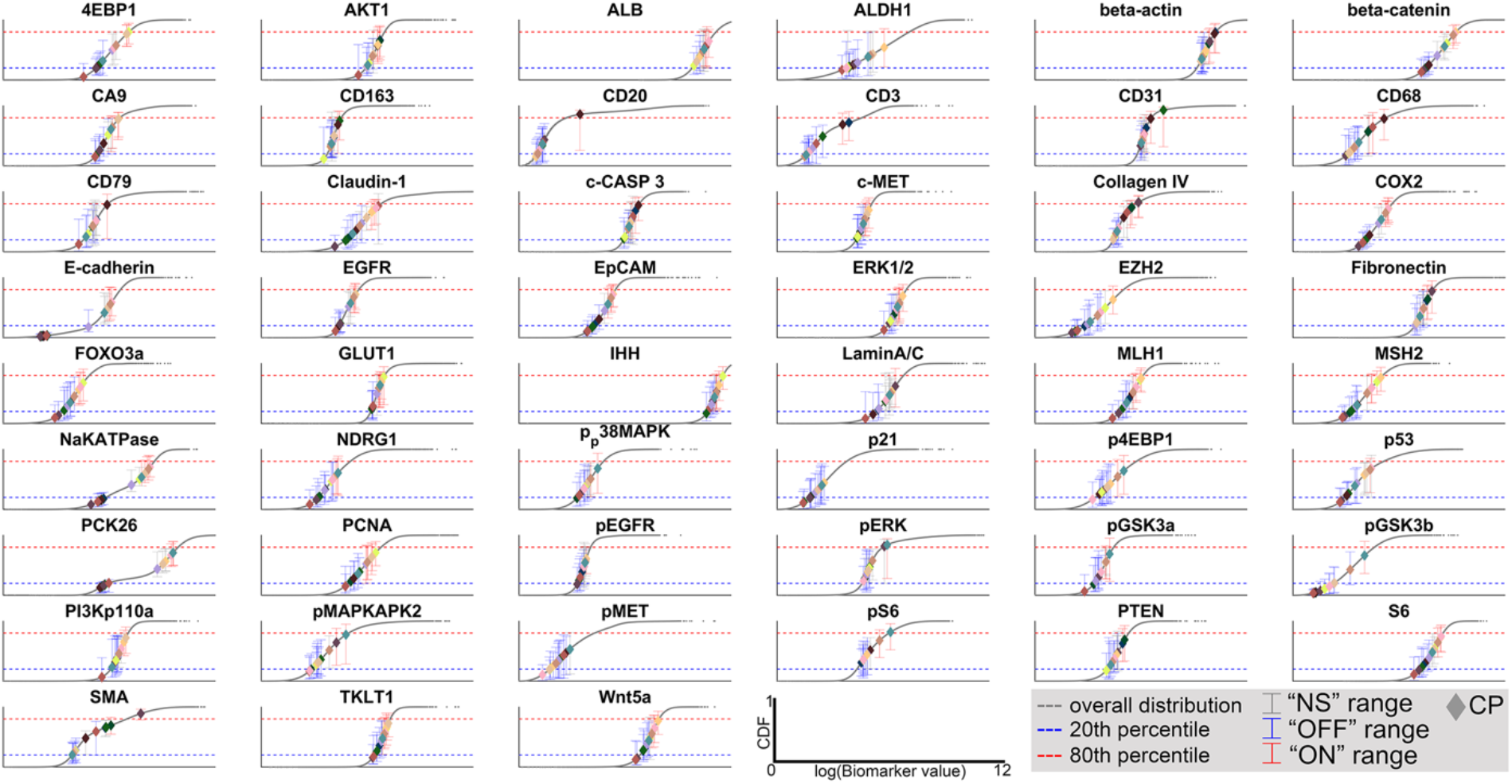
CP-specific biomarker classifications. Each CP is characterized with a unique biomarker signature (Figure 3b). The signatures are obtained by comparing the histogram of biomarker values over all cells (Figure S4) vs the histogram of biomarker values over the specialized cells in each CP (see Methods). The figure panel shows the cumulative distribution functions (CDFs) for all biomarkers (gray lines) and the 20^th^ and 80^th^ percentiles (see legend) in each CDF. The biomarker distribution over all specialized cells in each CP is represented by the mean (diamonds color-coordinated with Figure 4) and an error bar showing the 20^th^ and 80^th^ percentile values. Each biomarker is classified as “ON”, “OFF”, “NS” (see Methods for more details).

**Figure S8.**
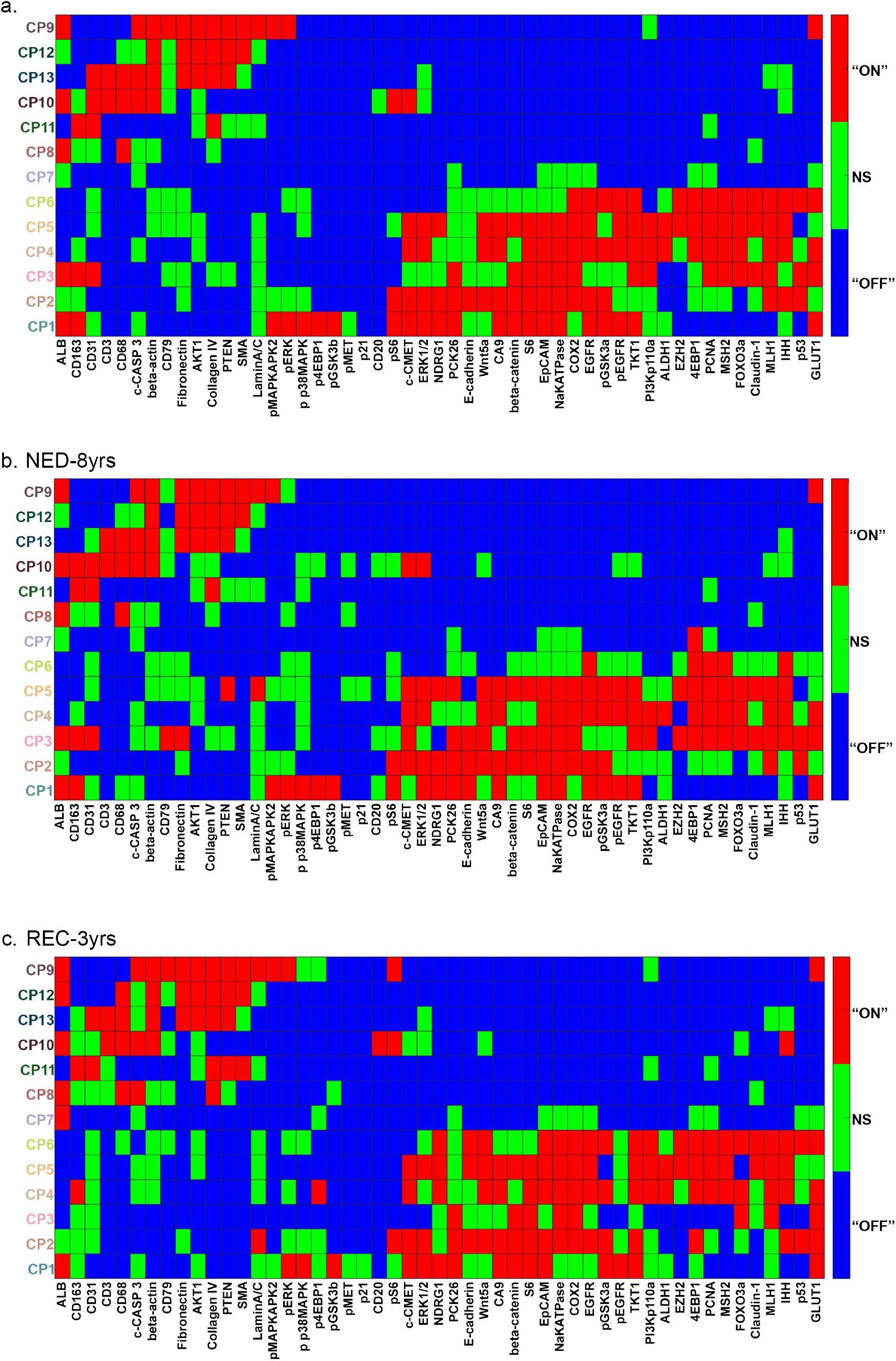
caption. The biomarker signatures are computed to facilitate qualitative interpretation only. The signatures shown in Figure 3b is an interpretation of each CP as a constellation of biomarkers based on the total CRC cohort of 213 patients. After reducing the patient cohort to the NED-8yrs and REC-3yrs sub-groups (91 patients), we re-interpret the CPs by computing the biomarker signatures (see Methods) based on only the reduced patient cohort (a). Further, we compute the CP biomarker signatures separately for the (b) NED-3yrs (45 patients) and (c) REC-8yrs (46 patients) cohorts. To compute the cohort-specific biomarker signatures, we are specifically comparing the biomarker distributions of each CP within each cohort compared to the biomarker distribution based on all cells in the reduced patient cohort (91 patients).

**Figure S9 –.**
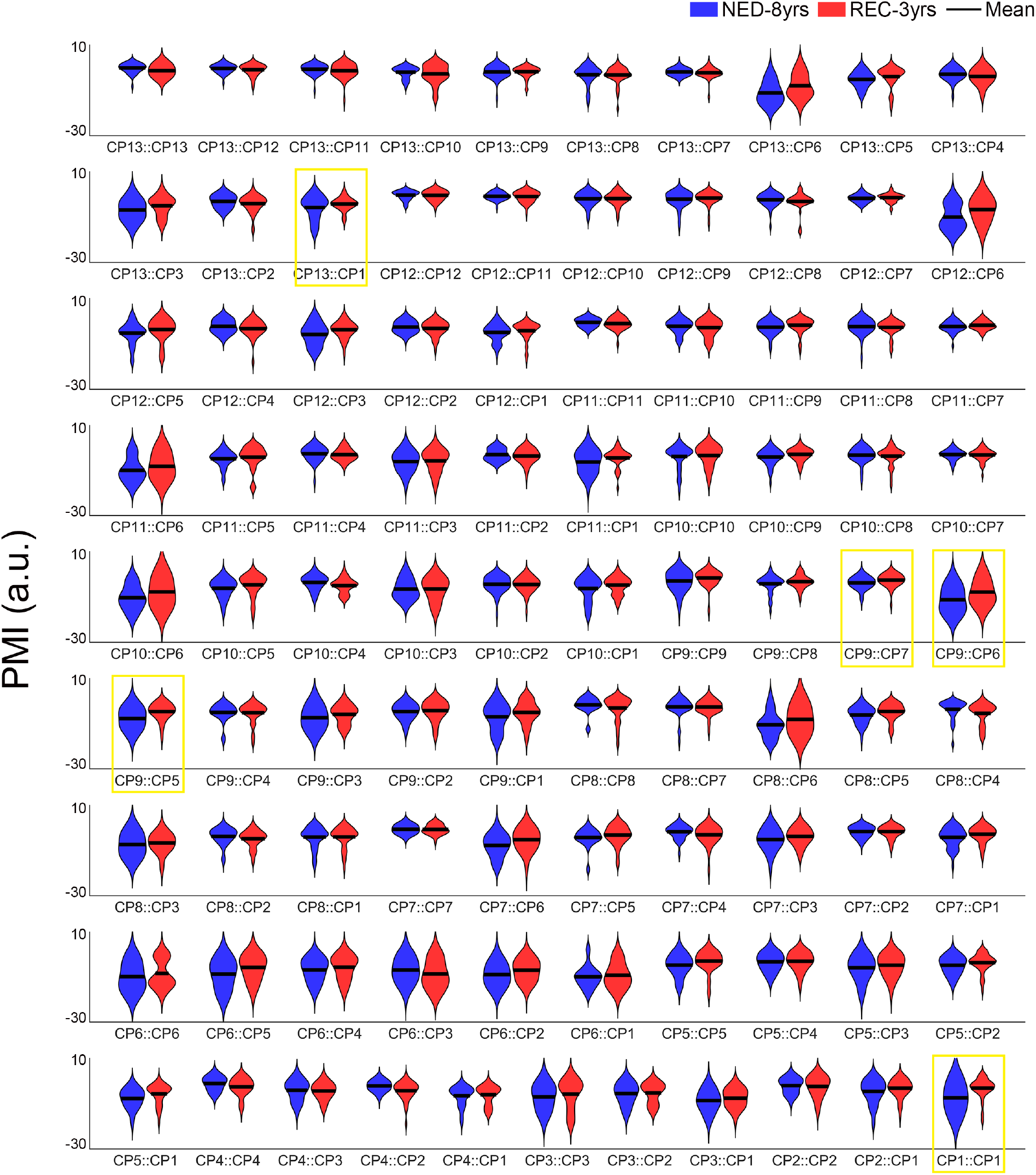
Distribution of PMI values. The distributions of PMI values for each CP-pair grouped by the outcome data (NED-8yrs in blue, REC-3yrs in red). Comparing the distributions using the Kendall rank correlation coefficients and the Wilcoxon rank sum test, we found 5 significant CP-pairs highlighted in yellow (see Methods and Table S4).

**Figure S10 –.**
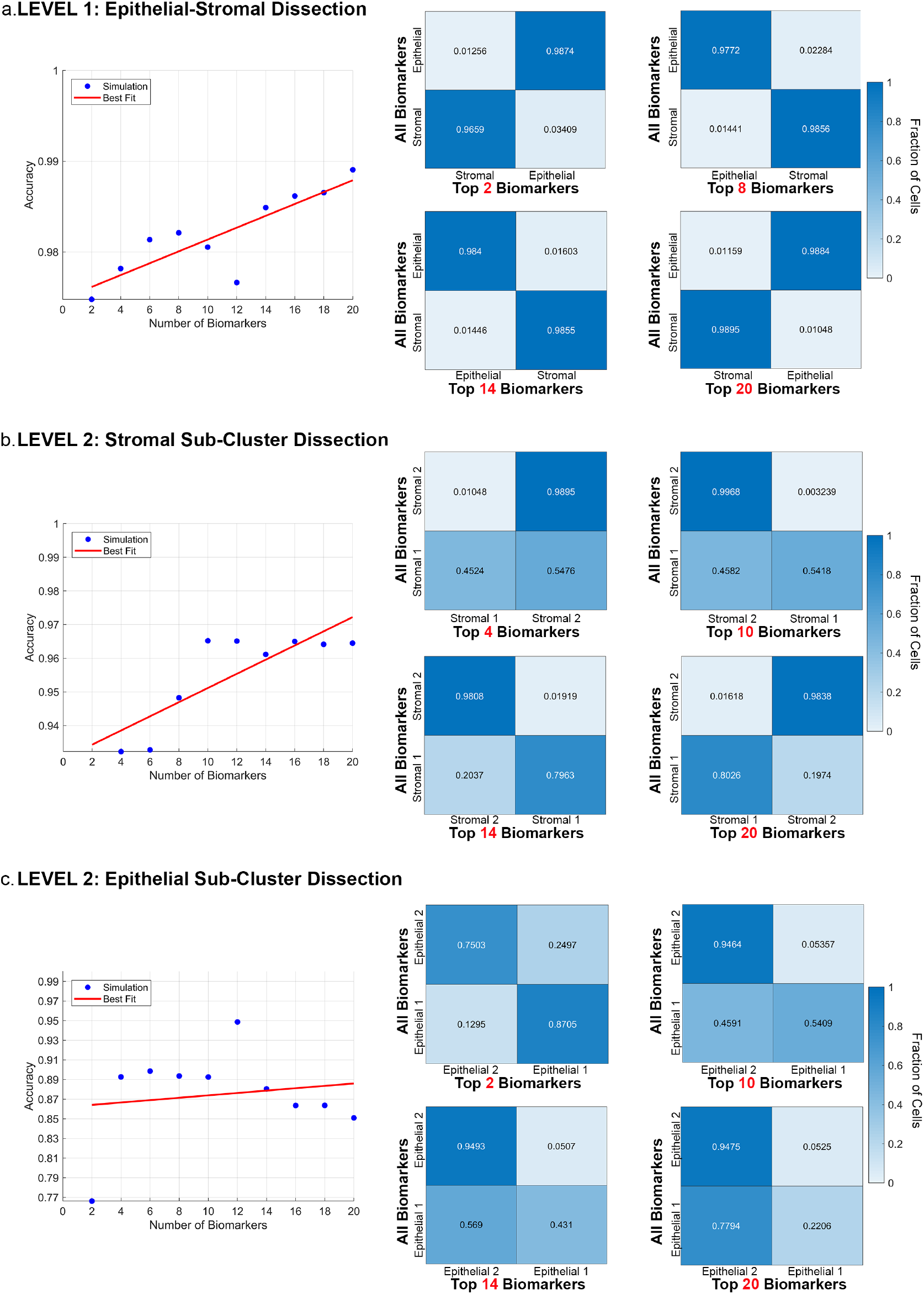
What is the minimal set of biomarkers needed to reproduce the LEAPH results with 51 biomarkers? The cyclical process of labeling, imaging, and quenching of the hyperplexing imaging platforms permits selecting biomarkers on demand. Because LEAPH determines an ordered list of key biomarkers at each level of the hierarchy, we performed a virtual simulation where the tissue is assumed to be labeled with a subset of the key biomarkers (as opposed to the 51 biomarkers that we started with). The key biomarkers are selected based on the order of discriminative biomarkers from the ALL-DATA results with 51 biomarkers (Table S7). LEAPH is then run to determine if the phenotype identities for each cell changed from the ones obtained with 51 biomarkers. For comparing phenotyping identities, we use the maximum CP ownership probability from the ALL-DATA results with 51 biomarkers (Figure 3) as the ground truth label for each cell. We measure the accuracy as the percentage of cells with a matching CP identity to the ALL-DATA results with 51 biomarkers. We ran this experiment to test the phenotypic identities at first two levels of the hierarchy: epithelial and stromal division at the first level and their individual subtypes at the second. **a.** As expected, the epithelial-stromal dissection is low-dimensional, only requiring 2 biomarkers (pck26, E-cadherin) to obtain an accuracy above 97%. Increasing the number of biomarkers further increases the accuracy to almost 99%. **b-c.** Both subtype dissections require a larger set of biomarkers than the first level. The stromal subtype dissection (**b**) reaches above 93% accuracy with 4 biomarkers (Lamin A/C, Claudin 1, Akt, 4EBP1). The addition of more biomarkers leads to an accuracy above 97% with 8 or more biomarkers. The epithelial subtype dissection (**c**) reaches 89% accuracy with 4 biomarkers (4EBP1, pS6, pMAPKAPK2, PCNA) but an increase in the number of biomarkers does not make a substantial difference in the reproduction accuracy.

## Notes

### Summary of Updates

Additional biological insights have bene added to the manuscript to make it complete. In support of this Fig 4d has been updated. A new supplementary figure has also been added.

